# RGI-Toolkit: Differentiable Restraints for Controllable Biomolecular Structure Prediction

**DOI:** 10.64898/2026.10.05.756905

**Authors:** Tatsuki Hori, Yoshitaka Moriwaki, Ryuichiro Ishitani

**Affiliations:** Department of Computational Drug Discovery and Design, Medical Research Laboratory, Institute of Integrated Research, Institute of Science Tokyo, 1-5-45 Yushima, Bunkyo-ku, Tokyo 113-8510, Japan; Graduate School of Agricultural and Life Sciences, The University of Tokyo, 1-1-1 Yayoi, Bunkyo-ku, Tokyo 113-8657, Japan; Department of Biological Sciences, Graduate School of Science, The University of Tokyo, 7-3-1 Hongo, Bunkyo-ku, Tokyo 113-0033, Japan

## Abstract

Diffusion-based models have substantially advanced biomolecular structure prediction. Within this framework, restraint-guided inference (RGI) corrects the denoiser output at each reverse-diffusion step using a differentiable restraint loss, allowing available experimental information, stereochemical requirements, or a specified target conformational state to be incorporated into predictions without retraining the model. Here, we present RGI-Toolkit, a general-purpose Python library that makes RGI available for a range of structure predictors. The toolkit consolidates restraint definition, atom selection, loss evaluation, and coordinate optimization into a shared, backend-agnostic engine, limiting predictor-specific code to a thin preprocessing adapter and a single hook in the denoising loop. Restraints on distances, angles, and dihedrals, on deviations from reference structures, and on user-defined reaction coordinates, together with ligand conformer restraints, are all specified in a common configuration format. The toolkit currently supports six predictors, including AlphaFold3. Across all six predictors, conformer restraints eliminated chirality and cis/trans errors in the predicted ligands and improved their local geometry, while geometric and custom restraints systematically shifted the sampled protein conformations toward the specified target states. RGI-Toolkit thus provides a unified framework for integrating experimental information and prior geometric knowledge into structure prediction and for generating biomolecular structures with specified conformational and stereochemical properties.

## Introduction

Diffusion-model-based structure predictors such as AlphaFold3 (AF3)^1^, Boltz^2,3^, Protenix^4^, Chai^5^, OpenFold3^6^, and ESMFold2^7^ have driven rapid advances in predicting the structures of complexes containing proteins, nucleic acids, and small molecules. However, it remains difficult to ensure that the predicted structures consistently reflect available experimental information, known partial structures, a desired domain arrangement, or chemically valid ligand geometry. Empirical strategies such as subsampling the input multiple sequence alignment (MSA) can broaden the range of sampled conformational states; however, they offer no direct means of steering predictions toward a desired state^8^.

We previously developed restraint-guided inference (RGI), a method that directly steers the inference process without modifying the trained model itself^9,10^. In AF3-type models, structures are generated through successive reverse diffusion steps, with a trained denoiser predicting a noise-free structure at each step to guide the update of atomic coordinates. We showed that structure prediction can be guided by arbitrary differentiable restraints imposed during this reverse diffusion process, without retraining the model. We implemented this framework in Boltz^2,3^ and applied it to eliminate stereochemical errors in small-molecule ligands^9^ and to sample protein conformational states^10^.

In the present work, we present RGI-Toolkit, a general-purpose toolkit that extends the applicability of RGI to structure predictors beyond Boltz, including AF3. RGI-Toolkit consolidates restraint definition, atom selection, energy evaluation, and coordinate optimization into a shared engine, confining predictor-specific modifications to a thin adapter in the preprocessing stage and a single hook in the inference loop. This design allows the same restraint configuration to be readily reused across different structure predictors and enables structure generation to be guided by combinations of ligand-geometry restraints, reference structures, and user-defined reaction coordinates. RGI-Toolkit has also been integrated into ColabFold^11,12^, enabling restraint-guided structure prediction to be performed directly in a web browser without local installation.

### Software Design and Implementation

#### Overview of restraint-guided inference

We first outline the restraint-guided inference (RGI) method^9^. RGI corrects the output of the denoiser *D*_*θ*_ of a pretrained diffusion model to reduce restraint violations and uses the corrected prediction in the subsequent coordinate update at each step. Let *z* ∈ ℝ^3*N*^ denote the atomic coordinates of a system containing *N* atoms, and let *L*_restr_(*z*) be a non-negative loss that measures restraint violations. The set of structures that satisfy the restraints is then defined as

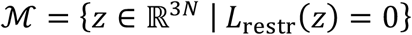

In RGI, this loss is minimized locally, starting from the structure *y*_*i*_ predicted by the denoiser at noise level *σ*_*i*_:

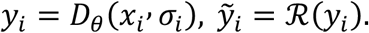

Here, *x*_*i*_ denotes the noisy atomic coordinates at step *i*, and *ℛ* denotes the local minimization procedure initialized at *y*_*i*_; if the loss converges to zero, then 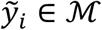. In this sense, the correction can be interpreted as a generalized projection onto the restraint set *ℳ*. The corrected prediction is then used for the next coordinate update. For the Euler-type update in the EDM framework^13^, omitting the additional noise injection and setting the step scaling factor to unity yields

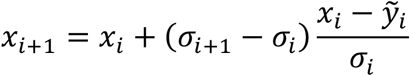

RGI thus modifies the update direction of the generative process by correcting the denoised prediction according to the restraints. This provides an approximate means of generating structures that satisfy the restraints without retraining the model. Full details of the RGI method are given in ref^9^. RGI-Toolkit provides a flexible interface for defining the loss *L*_restr_(*z*) as an arbitrary differentiable function, together with a general framework for performing the local minimization *ℛ* using gradient-based optimizers.

#### Toolkit architecture

The implementation consists of four components: (i) preprocessing adapters that convert atom, residue, and ligand information into a common representation; (ii) a backend-agnostic shared engine layer that builds restraint specifications; (iii) an optimizer layer that evaluates restraint losses, computes their gradients using automatic differentiation in PyTorch^14^ or JAX^15^, and optimizes atomic coordinates; and (iv) predictor hooks that are inserted into the denoising loop of each structure prediction backend (Fig. 1A). Six models are currently supported: Boltz-2, Protenix-v2, AlphaFold3, Chai-1, OpenFold3, and ESMFold2. These models employ similar EDM-based sampling schemes for structure generation, allowing the RGI predictor hook to be inserted immediately after each denoiser call to correct the predicted coordinates before the next coordinate update (Fig. 1A). The shared engine supports both L-BFGS minimization^16^ and a nonlinear conjugate gradient method^17^ implemented within RGI-Toolkit.

**Figure 1.**
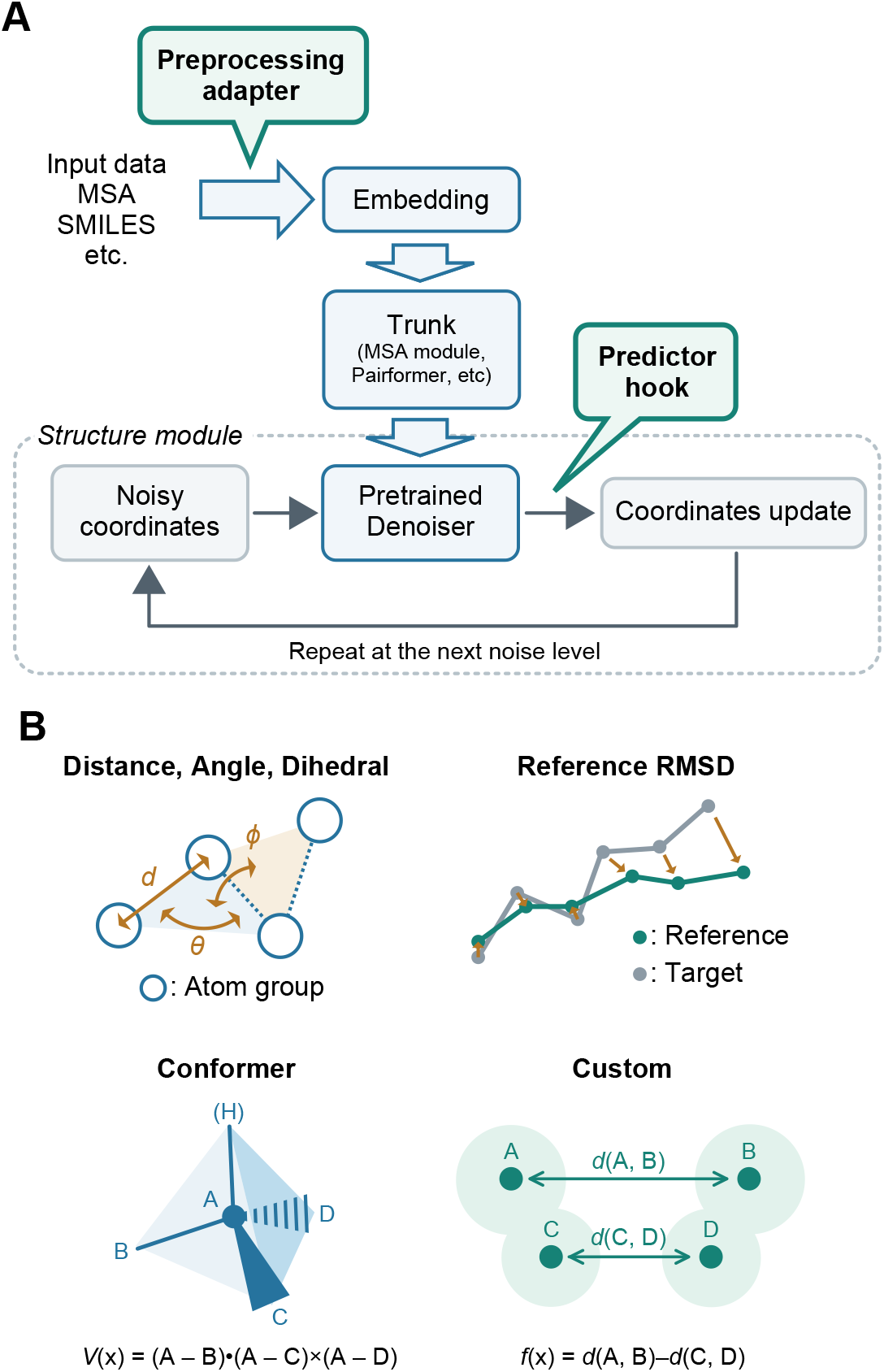
Overview of RGI-Toolkit. (A) Restraint-guided reverse diffusion in an AF3-type structure predictor. A preprocessing adapter and a predictor hook (green) are the only predictor-specific components; the hook corrects the denoised coordinates according to the restraints before each coordinate update. (B) Main restraint types: distance (*d*), angle (*θ*), and dihedral (*ϕ*) between the geometric centroids of atom groups; RMSD from a reference structure after optimal superposition; conformer restraints, which comprise several terms enforcing consistency with an ideal conformer (the chirality term, based on the signed volume *V*(*x*), is shown as an example); and custom restraints *f*(*x*) defined by user-specified expressions, such as the difference between two distances.

#### Supported restraint types

RGI-Toolkit provides the following types of restraints (Fig. 1B).

##### Distance, angle, and dihedral

restraints that control the distance, angle, or dihedral angle defined by the geometric centroids of two to four atom groups, using either a harmonic or a flat-bottom potential (Fig. 1B).

##### Reference RMSD

restraints that drive the root-mean-square deviation (RMSD) of selected atoms from a reference structure toward a specified target value after optimal superposition using the Kabsch algorithm^18^ (Fig. 1B). Restraints to multiple reference structures can be applied simultaneously. Atom groups in an external PDB or mmCIF file can also be used as fixed reference points, allowing selected parts of the predicted structure to be guided toward known coordinates.

##### Conformer

restraints that promote stereochemical consistency in the predicted structure using an ideal conformer as a reference. Individual terms are available for bond lengths (bond), bond angles (angle), chirality (chiral), cis/trans isomerism (cistrans), and intermolecular (i.e., protein-ligand) van der Waals repulsion (vdw), with independently adjustable weights. The cistrans term restrains dihedral angles about acyclic double bonds. The ideal conformer is generated using the ETKDGv3 algorithm^19^ implemented in RDKit and optimized with the universal force field^20^.

##### Custom

restraints defined by mathematical expressions constructed from quantities such as the geometric centroids of selected atom groups and the distances between them (Fig. 1B). These expressions are evaluated by a parser, allowing users to define low-dimensional reaction coordinates. Gradients of the specified expressions are computed by automatic differentiation using the deep learning framework employed by the structure prediction model.

These restraints can be used individually or in combination.

#### Restraint configuration

Restraints are specified in YAML or JSON format, or as a Python dictionary for ESMFold2. Atom selections are defined using a domain-specific language that combines chain names, residue numbers, atom names, and logical operators. For distance, angle, and dihedral restraints, the geometric centroids of selected atom groups can serve as the points defining the restrained geometry. The configuration format is backend-agnostic, allowing the same restraint definition to be reused across different structure prediction models. The implementation details of RGI-Toolkit are provided in the Supplementary Information.

### Applications and Validation

#### Benchmark design

Two benchmarks were conducted: a ligand conformer benchmark to evaluate the effectiveness of ligand conformer restraints and a conformational state-control benchmark to assess the ability of distance, angle, reference RMSD, and custom restraints to steer proteins toward specified conformational states. Six structure prediction models were evaluated: Boltz-2, Protenix-v2, AlphaFold3, Chai-1, OpenFold3, and ESMFold2.

For the ligand conformer benchmark, we selected 209 protein–ligand complexes whose ligands contain double bonds with distinguishable *cis*/*trans* isomerism from the PLINDER^21^-based benchmark dataset used by Ishitani et al.^9^. Bond lengths, bond angles, chirality, *cis*/*trans* isomerism, and steric clashes with the protein were assessed using the RDKit-based evaluation protocol^9^ and PoseBusters^22^. For each structure prediction model, predictions generated with the ligand conformer restraints of RGI-Toolkit were compared with unrestrained predictions. A single prediction with one random seed was generated for each complex under each condition. For Boltz-2 and Protenix-v2, the ligand geometry guidance methods native to those implementations^3,23^ were also included in the comparison.

The conformational state-control benchmark examined transitions between the open and closed states of glutamine-binding protein (QBP)^24,25^ and adenylate kinase (ADK)^26,27^, and between the outward-facing and inward-facing states of the major facilitator superfamily (MFS) transporter DgoT^28^. For each protein, a common MSA generated with ColabFold^11^ was used across all six structure prediction models. In the RGI condition, two separate benchmarks were performed for QBP, one with an interdomain distance restraint and the other with reference RMSD restraints to both the open and closed structures. ADK was subjected to an interdomain angle restraint, and DgoT to a custom restraint on the difference between two interdomain distances. For each benchmark case, 45 structures per model were generated under the RGI condition, using five target values of the reaction coordinate and nine random seeds per target value. Predictions under the baseline condition (unrestrained prediction with the full MSA) and the MSA subsampling condition were generated following the protocols used in Hori et al.^10^. Predicted structures were evaluated using TM-scores relative to the experimental structures of the open and closed states for QBP and ADK, and of the inward-facing and outward-facing states for DgoT. For all restraints, the values of the restrained quantities were also calculated from the predicted structures and compared with their target values.

#### Improvement in ligand stereochemistry and geometry

Under the baseline condition without restraints or guidance, the frequency of stereochemical errors varied across structure prediction models (Fig. S1). The chirality agreement rate with the input molecules was around 50% across all models, and the *cis*/*trans* agreement rate ranged from 70% to 80% across all models. In contrast, with RGI conformer restraints, the agreement rates for both chirality and *cis*/*trans* isomerism reached 100% for all six models (Fig. S1A, B). The median bond length and bond angle RMSDs relative to the ideal conformer were lower than those under the baseline condition for every model (Fig. S1C, D). The native steering method of Boltz-2 also improved the chirality agreement rate to 100% and the cis/trans agreement rate to 99%. Similarly, the training-free guidance native to Protenix-v2 improved the chirality agreement rate to 98.6% and the *cis*/*trans* agreement rate to 82.3%. However, both native methods yielded higher median bond length RMSDs than RGI. The minimum relative ligand–protein distance was also improved by the van der Waals repulsion term, and with RGI it was kept above the specified lower bound (*α* = 0.75) for all models (Fig. S1E). Consistent with our previous study^9^, however, RGI neither improved nor degraded the accuracy of ligand pose prediction (Fig. S1F). Figs. S2-S5 compare the restraint target values with the values achieved in the predicted structures for the chirality, cis/trans, bond-length, and bond-angle terms. The restraints were well satisfied not only for chirality and cis/trans isomerism but also for bond lengths and bond angles. These results demonstrate that RGI improves ligand stereochemistry consistently, bringing all six structure prediction models to comparable levels of accuracy. Further results of the ligand conformer benchmark are discussed in the Supplementary Information.

#### Controlled sampling of protein conformational states

For QBP, we applied an interdomain distance restraint and increased the target distance from 25 Å, the value in the experimental closed structure, to 29.1 Å, the value in the experimental open structure. As expected, increasing the target distance progressively decreased the TM-score relative to the closed reference and increased the TM-score relative to the open reference for all models (Fig. S6). The interdomain distances measured in the predicted structures closely matched the target values in all models, indicating that the distance restraint was almost fully satisfied (Fig. S7). For ADK, applying an interdomain angle restraint and varying the target angle from 60.9° to 84.8° (the values in the experimental closed and open structures, respectively) yielded a continuous progression from closed-like to open-like conformations for all models (Fig. S8). The measured interdomain angles closely followed the target values in all models, with small deviations (Fig. S9).

To control the conformational state of QBP using RMSD restraints, we applied two reference RMSD restraints simultaneously. Denoting the RMSD between the open and closed reference structures by *T* (*T* = 5.3 Å for QBP), we set the target RMSD to the open reference to *t* and that to the closed reference to *T* − *t*. As *t* was varied, the two target RMSD values changed linearly in opposite directions, and predictions were generated at five values of *t* from 0 to *T*. For all models, the mean TM-score varied systematically with the target values, and the TM-scores at the two endpoints agreed well with those of the experimental reference structures across models (Fig. S10). The RMSD values of the predicted structures to the open and closed references did not exactly match the target values *t* and *T* − *t* (Fig. S11). These residual deviations are inherent to the restraint design rather than a failure of the restraints, because the triangle inequality gives *r*_open_ + *r*_closed_ ≥ *T*, with equality only for structures lying on the linear interpolation between the two references. Indeed, the predicted structures were reasonable in terms of TM-scores, stereochemical quality, and visual inspection, and the results closely matched those obtained with the interdomain distance restraint (Fig. S6), indicating that, for QBP, distance and RMSD restraints enable comparable sampling of conformational states.

To control the conformational state of DgoT during its transport cycle, we defined a custom restraint on the difference *ΔD* = *D*_in_ − *D*_out_ between the cytoplasmic gate distance *D*_in_ = *d*(A,B) and the periplasmic gate distance *D*_out_ = *d*(C,D). Here, A and B denote groups of transmembrane helices on the cytoplasmic side, and C and D denote those on the periplasmic side (the atom selections are given in the Supplementary Information). Varying the target value of *ΔD* from −4.3 to 5.9 Å enabled all models to sample a series of structures that included conformations resembling the experimental outward- and inward-facing structures (Fig. S12). Further results of the conformational state-control benchmark are discussed in the Supplementary Information.

## Limitations

RGI-Toolkit is a framework that helps diffusion-based structure prediction models toward structures that satisfy user-specified restraints; it does not guarantee that the restraints themselves are physically or biologically reasonable. Excessively large restraint weights, mutually inconsistent restraints, and incomplete atom correspondence between the predicted and reference structures in RMSD restraints can introduce local strain or lead to sampling failures. Results should therefore be evaluated carefully: predictions should be run with multiple random seeds and compared with unrestrained controls, and prediction confidence scores (e.g., pLDDT), stereochemical quality (e.g., assessed with MolProbity^29^ and PoseBusters^22^), and restraint residuals should be examined. Relaxing RGI-generated structures using molecular dynamics simulations, followed by resampling, is also an effective strategy^10^. The effectiveness of conformer restraints depends on the quality of the reference coordinates provided for the input molecule. If 3D embedding produces erroneous reference coordinates, as can happen with the ETKDG algorithm in RDKit, the resulting restraints may fail to guide sampling toward the correct structure. Iterative optimization at each reverse-diffusion step introduces additional computational overhead, which increases with the number of optimization iterations, the number of selected atoms, and the number of restraints.

## Conclusion

RGI-Toolkit provides a unified framework for incorporating differentiable restraints into the denoising loops of diffusion-based biomolecular structure prediction models. A shared restraint engine, together with thin model-specific adapters and hooks, allows a single configuration scheme to be applied across six structure prediction models without modifying their weights or training procedures. In the ligand conformer benchmark, conformer restraints consistently improved ligand stereochemistry and local geometry across all six models. In the protein state-control benchmark, the sampled conformations shifted progressively as the target values for distances, angles, RMSDs relative to reference structures, and user-defined reaction coordinates were varied. Beyond the restraint types evaluated in these benchmarks, RGI-Toolkit also supports dihedral restraints defined over atom groups, and restraints that anchor selected atoms to an external reference structure. Taken together, RGI-Toolkit provides a practical means of incorporating experimental data and other prior knowledge into structure prediction, generating specific conformational states for hypothesis testing, and systematically comparing predictions across models.

## Data and Software Availability

The software will be available under the MIT License at https://github.com/cddlab/rgi_toolkit. The MIT-licensed core RGI-Toolkit library is available for both academic and commercial use. In addition, an RGI-enabled version of ColabFold is available as a Google Colab notebook at https://github.com/sokrypton/ColabFold/blob/main/ColabFold2_preview.ipynb, which allows users to run restraint-guided structure prediction without local installation. The PLINDER-derived benchmark dataset used in this study (plinder-cistrans-209) is available on Hugging Face at https://huggingface.co/datasets/wasarou/plinder-cistrans-209.

## Author contributions

T.H. designed the study, performed computational experiments, analyzed the data and wrote the manuscript. R.I. and Y.M. analyzed the data and wrote the manuscript. R.I. directed and supervised this study.

## Acknowledgements

This work was supported in part by JST BOOST (JPMJBS2430 to T.H.); the Platform Project for Supporting Drug Discovery and Life Science Research (BINDS) from AMED (JP25ama121012 to R.I.); JSPS KAKENHI Grant-in-Aid for Transformative Research Areas (A) (25H01570 and 25H02250 to R.I.) and for Scientific Research (25K01890 to Y.M.); the Medical Research Center Initiative for High Depth Omics at Institute of Science Tokyo; the Joint Usage/Research Center for intractable diseases (Nanken-Kyoten 2026) at Institute of Science Tokyo; and the Multi-layered Stress Diseases Project at Institute of Science Tokyo (JPMXP1323015483).

