## Supplementary material for "RGI-Toolkit: Differentiable Restraints for Controllable Biomolecular Structure Prediction": suppl

Tatsuki Hori[a]

Yoshitaka Moriwaki[a,b]

Ryuichiro Ishitani[a,c,*]

[a]*Department of Computational Drug Discovery and Design, Medical Research Laboratory, Institute of Integrated Research, Institute of Science Tokyo, 1-5-45 Yushima, Bunkyo-ku, Tokyo 113-8510, Japan;* [b]*Graduate School of Agricultural and Life Sciences, The University of Tokyo, 1-1-1 Yayoi, Bunkyo-ku, Tokyo 113-8657, Japan;* [c]*Department of Biological Sciences, Graduate School of Science, The University of Tokyo, 7-3-1 Hongo, Bunkyo-ku, Tokyo 113-0033, Japan*

* Corresponding author

### Supplementary Text

#### Implementation details

The ligand conformer restraints used in this work extend the previously reported bond-length, bond-angle, and chirality restraints[1] with two additional terms: a *cis*/*trans* restraint and a protein-ligand van der Waals repulsion restraint. The *cis*/*trans* restraint applies to acyclic double bonds for which a dihedral angle can be defined. It is implemented as a harmonic restraint on the dihedral angle defined by the coordinates of the four atoms comprising the double bond and one substituent on each side, with a target value taken from the reference conformer. The van der Waals repulsion restraint imposes a lower bound on interatomic distance of each nonbonded protein-ligand atom pair in a specified set $P$:

$$L_{\mathrm{vdW}} = \sum_{(i,j)\in P} \min\left(0, d_{ij} - \alpha\,(R_i + R_j)\right)^2$$

Here, $d_{ij}$ is the distance between atoms $i$ and $j$, $R_i$ and $R_j$ are their respective van der Waals radii obtained from the RDKit periodic table, and $\alpha$ is a scaling factor. The total loss function for the ligand conformer restraints is given by

$$L_{\mathrm{ligand}} = w_b L_{\mathrm{bond}} + w_a L_{\mathrm{angle}} + w_c L_{\mathrm{chiral}} + w_t L_{\mathrm{cistrans}} + w_v L_{\mathrm{vdW}}.$$

where $w_b$, $w_a$, $w_c$, $w_t$, and $w_v$ are the weights of the individual terms.

For the restraints used to control conformational states, let $x_i$ denote the coordinates of atom $i$ belonging to atom group $G_k$ ($k = 1, \ldots, 4$). The unweighted mean of the atomic coordinates,

$$c_k = \frac{1}{\mid G_k \mid} \sum_{i \in G_k} x_i,$$

defines the geometric centroid of the group and serves as the reference point for evaluating the restrained quantities. The distance restraint acts on the distance between the geometric centroids of two atom groups, $d(G_1, G_2) = \|c_2 - c_1\|$, and drives this distance toward a specified target value. Similarly, the angle restraint acts on the angle formed by the centroids of $G_1$, $G_2$, and $G_3$, and the dihedral restraint acts on the dihedral angle formed by the centroids of $G_1$ through $G_4$; each quantity is driven toward its specified target value.

For the reference RMSD restraint, the reference structure is superposed onto the predicted structure by the Kabsch algorithm[2] using the set of atoms designated as fit atoms. The RMSD is then evaluated over a separately specified set of calculation (calc) atoms. For reference structure $s$, the measured quantity $r_s(x)$ is defined as

$$r_s(x) = \sqrt{\frac{1}{n_s} \sum_{a \in C_s} \left\| x_a - q_{s,a} \right\|^2},$$

where $q_s$ denotes the coordinates of the reference structure after superposition onto the predicted structure $x$, $C_s$ is the set of corresponding calc atoms, and $n_s = |C_s|$. The Kabsch superposition is recomputed from the denoised coordinates at every reverse diffusion step and is also updated at every iteration of the subsequent minimization loop.

The loss is computed from the measured quantities as follows. For each restraint $i$, the deviation $\delta_i$ of the measured value $r_i$ from its target value is computed, and the loss is defined as the weighted sum of the squared deviations with weights $w_i$:

$$L = \sum_i w_i \ \delta_i^2.$$

The definition of $\delta_i$, and hence the form of the loss function, can be selected from four options. For the harmonic form,

$$\delta_i = r_i - t_i;$$

for the flat-bottom form,

$$\delta_i = \min\left(0, \ r_i - t_{i,1}\right) + \max\left(0, \ r_i - t_{i,2}\right);$$

where $t_{i,1}$ and $t_{i,2}$ ($t_{i,1} \leq t_{i,2}$) are the lower and upper bounds, respectively; and for the flat-bottom1 and flat-bottom2 forms, respectively,

$$\delta_i = \min\left(0, \ r_i - t_{i,1}\right), \ \delta_i = \max\left(0, \ r_i - t_{i,2}\right).$$

The flat-bottom form yields zero loss within the interval $t_{i,1} \leq r_i \leq t_{i,2}$, whereas flat-bottom1 and flat-bottom2 impose only a lower bound and an upper bound, respectively. Only the harmonic form was used in the state-control benchmarks of this study. For custom restraints, a differentiable expression constructed from quantities such as the inter-centroid distances defined above is evaluated directly as the loss function. The loss function was minimized using the conjugate gradient (CG) method, with the maximum number of iterations specified by the parameter max_iter.

The range of reverse diffusion steps over which each restraint is active is specified in terms of the noise level $\sigma$: a restraint is applied only when $\sigma_{\text{stop}} \leq \sigma \leq \sigma_{\text{start}}$, where $\sigma_{\text{start}}$ and $\sigma_{\text{stop}}$ are set by the parameters start_sigma and stop_sigma, respectively. If start_sigma is omitted, the restraint is applied from the first step; if stop_sigma is omitted, the restraint remains active until the final step.

#### Conformer benchmark details

In the ligand conformer restraint benchmark, the conformer restraints were applied to the ligands in the 209-entry *cis*/*trans* benchmark set. For each entry, a single input MSA in a3m format, generated with ColabFold[3], was shared across all six models. All conformer restraint settings were left at their default values except for start_sigma, which was set to 1.0; stop_sigma remained unset. The conformer restraints were therefore activated once the noise level decreased to $\sigma = 1.0$ and remained active until the end of the reverse-diffusion process. Thus, the restraints acted only during the later stages of reverse diffusion, after the overall fold had been established. The scaling factor $\alpha$ of the van der Waals repulsion term was 0.75, and the maximum number of iterations in the CG minimization (max_iter) was the default value of 100.

The target values for the bond, angle, chiral, and cistrans terms were derived from the ligand reference conformer provided by each predictor implementation after further energy minimization with the Universal Force Field (UFF)[4]. To verify that this local relaxation did not alter the stereochemistry of the reference conformer, tetrahedral chirality and distinguishable *cis*/*trans* configurations were reassigned from the relaxed three-dimensional coordinates and compared with the corresponding CIP labels defined on the molecular graph. If any mismatch was detected, a new conformer was generated using ETKDGv3[5] with a different deterministic seed, and the same local relaxation and stereochemical validation steps were repeated, up to a maximum of four attempts.

In the ligand conformer benchmark, a single prediction (one random seed) was generated for each entry under each condition. Ligand predictions were obtained for all 209 entries under every condition with all six models, so all entries were used for the comparisons. All metrics were defined per entry rather than per atom. Stereochemistry was assessed by comparing the tetrahedral chirality and the *cis*/*trans* configurations of acyclic double bonds in the predicted ligand with those of the input molecule constructed with RDKit. For each entry, match was set to 1 if all stereocenters targeted by the RGI restraints agreed with those of the input molecule and to 0 otherwise. Likewise, ez_match was set to 1 if all distinguishable *cis*/*trans* double bonds agreed with those of the input molecule and to 0 otherwise. Bond-length and bond-angle RMSDs were calculated relative to the corresponding ideal values obtained from a reference conformer prepared as described above. The ligand-protein minimum relative distance was defined as the smallest value of $d_{ij}/(R_i + R_j)$ over all ligand–protein atom pairs, where $d_{ij}$ is the distance between atoms $i$ and $j$ and $R_i$ and $R_j$ are their element-based van der Waals radii. Ligand RMSD was calculated as described previously[1]. All of these continuous metrics are reported as medians across entries.

#### Conformational-state control benchmark details

In the state-control benchmarks (distance, angle, RMSD, and custom restraints), harmonic restraints were used throughout. Unless otherwise specified, start_sigma and stop_sigma were left unset, so that the restraints were active from the first to the final step of the reverse-diffusion process, and the maximum number of iterations in the CG minimization (max_iter) was set to 100. Predictions were generated using nine random seeds for each target value. The baseline condition (unrestrained predictions using the full MSA) and the MSA-subsampling condition served as controls; predictions under these conditions were generated following the protocol described by Hori et al.[6]. Residue numbers given below are sequential indices starting at 1 within each chain.

The distance-restraint benchmark targeted glutamine-binding protein (QBP), using PDB entries 1GGG (open)[7] and 1WDN (closed)[8] as reference structures. The restraint was applied to the distance between the geometric centroids of two domains, the first comprising residues 4–83 and 185–223 and the second comprising residues 89–179.

The angle-restraint benchmark targeted adenylate kinase (ADK), using PDB entries 4AKE (open)[9] and 1AKE (closed)[10] as reference structures. The restraint was applied to the angle defined by the geometric centroids of three domains: the NMP domain (residues 30–59), the CORE domain at the vertex (residues 1–29, 60–121, and 160–214), and the LID domain (residues 122–159).

The RMSD-restraint benchmark used the same QBP system and reference structures as the distance-restraint benchmark, with restraints to both references applied simultaneously. Let $T$ denote the RMSD between the two reference structures after superposition, and let $q_{\text{open}}$ and $q_{\text{closed}}$ denote their respective coordinates after superposition onto the predicted structure. Following the notation above, $r_s(x) = \text{RMSD}(x, q_s)$ for $s$ = open, closed. For a target value $t$, the restraint loss function was defined as:

$$L_{\text{RMSD}} = \left(r_{\text{open}}(x) - t\right)^2 + \left(r_{\text{closed}}(x) - (T - t)\right)^2$$

Both the fit and calc atom selections were restricted to Cα atoms in the reference and predicted structures. Thus, the restraint directly adjusted only the selected Cα coordinates, while each predictor reconstructed the local geometry, including side chains. For this benchmark only, stop_sigma was set to 1.5 to deactivate the restraints during the final low-noise stage of the reverse-diffusion process. In addition, max_iter was increased to 1000 for this benchmark to allow both RMSD terms to be optimized simultaneously. The Cα RMSD between the open and closed references was 5.338 Å, and $T$ was set to 5.3 Å. Because the RMSD after optimal superposition satisfies the triangle inequality, $r_{\text{open}}$ +

$r_{\text{closed}} \geq T$ always holds, with equality only when the predicted coordinates lie on the linear interpolation between the optimally aligned coordinates of the two references. Because the actual domain motion of QBP involves a hinge rotation, intermediate conformations deviate from this linear interpolation and satisfy $r_{\text{open}} + r_{\text{closed}} > T$. Thus, the two target values $t$ and $T - t$ cannot be satisfied simultaneously. Furthermore, the RMSD restraints were removed in the late stage of reverse diffusion, allowing additional deviations from the target values. To assess whether these residual deviations affected the predicted conformational states, we evaluated the stereochemical quality of the predicted structures, as described below, and compared the predicted conformations with those obtained in the distance-restraint benchmark (see Supplementary Text).

The custom-restraint benchmark targeted the major facilitator superfamily (MFS) transporter DgoT, using PDB entries 6E9O (outward-facing) and 6E9N (inward-facing)[11] as reference structures. Four transmembrane helix groups were defined as follows: the cytoplasmic side of the N-terminal domain (A), residues 1–17, 52–66, 69–78, 108–121, 125–139, and 168–178; the cytoplasmic side of the C-terminal domain (B), residues 229–241, 280–295, 298–311, 341–354, 358–371, and 400–412; the periplasmic side of the N-terminal domain (C), residues 18–34, 38–51, 79–87, 92–107, 140–155, and 158–167; and the periplasmic side of the C-terminal domain (D), residues 242–250, 265–279, 312–323, 325–340, 372–386, and 388–399. Using the distances between the geometric centroids of these groups, the control variable was defined as $\Delta D = D_{\text{in}} - D_{\text{out}} = d(A, B) - d(C, D)$. The target value of $\Delta D$ was swept from −4.3 Å (outward-facing) to 5.9 Å (inward-facing).

The extent to which the predicted structures approached the intended conformational states was assessed using TM-scores, calculated with US-align[12] by independently aligning each predicted structure to each of the two reference structures.

In addition, prediction confidence was evaluated using the pLDDT and pTM scores reported by each model, and the stereochemical quality of the predicted structures was evaluated using MolProbity[13].

#### Detailed results of the ligand conformer benchmark

The median bond-length RMSD relative to the ideal conformer ranged from 0.0348 to 0.0560 Å across models under the baseline condition, compared with 0.0030 to 0.0043 Å under the RGI condition (Fig. S1C). Similarly, the median bond-angle RMSD ranged from 2.611 to 4.332° under the baseline condition but decreased to a range of 1.144 to 1.291° under the RGI condition (Fig. S1D). Even ESMFold2, which showed the largest bond-length and bond-angle RMSDs under the baseline condition, yielded values of 0.0031 Å and 1.165°, respectively, with RGI. Thus, RGI reduced both local geometric errors and their variation across models. We also compared individual bond lengths and bond angles in the predicted structures with the corresponding restraint target values (Figs. S2 and S3). These geometric parameters deviated substantially from the ideal values under the baseline condition, whereas with RGI they closely matched the ideal values, indicating improved local geometry.

The effect of the native guidance methods on local geometry varied with the model and the metric. Both native guidance methods improved the agreement rates for chirality and *cis*/*trans* configurations (Fig. S1A, B), but their effects on bond lengths and bond angles differed. The native steering of Boltz-2 increased the median bond-length RMSD from 0.0348 to 0.0419 Å and the median bond-angle RMSD from 2.611 to 3.826°

compared with the baseline (Fig. S1C, D). In contrast, the training-free guidance of Protenix-v2 decreased them from 0.0528 to 0.0324 Å and from 4.132 to 3.900°, respectively (Fig. S1C, D). These values for Protenix-v2 nevertheless remained larger than those obtained with RGI for Protenix-v2 (0.0035 Å and 1.194°, respectively; Fig. S1C, D). In the comparison of target and achieved values, some bond lengths and bond angles deviated more from their ideal values under native guidance than under the baseline condition for both Boltz-2 and Protenix-v2 (Figs. S2 and S3). This deterioration can be attributed to the different implementation of the native guidance potentials from that of RGI. These results indicate that, with native guidance, improved agreement in chirality and *cis*/*trans* configurations is not necessarily accompanied by a reduction in bond-length and bond-angle RMSDs.

For the chiral volumes, the comparison of target and achieved values showed that the sign of the chiral volume was inverted in about half of the cases under the baseline condition, whereas with RGI all of them were guided to the correct volumes (Fig. S4). The native guidance of Boltz-2 and Protenix-v2 also reduced sign inversions, but the deviations of the target chiral volumes were larger than those with RGI (Fig. S4). This difference may reflect the use of chirality corrections that do not explicitly restrain the chiral volume in either native guidance method. In addition to enforcing the correct handedness, a chiral-volume restraint constrains a geometric quantity that depends on the distances and angles between substituents around the stereocenter. It can therefore help maintain local geometry. By contrast, the methods adopted in the native guidance implementations can correct only the handedness of the stereocenter. For the cis/trans term, the comparison of target and achieved values likewise showed that RGI produced the correct configurations, in contrast to the baseline condition (Fig. S5). Native guidance also improved agreement in the cis/trans configurations, but because these methods do not strictly restrain the configuration, incorrect configurations remained in some cases (Fig. S5).

For the minimum relative distance between protein and ligand atoms, the RGI condition narrowed the distributions for all six models and suppressed the low-value tail observed under the baseline condition (Fig. S1E). With RGI, the minimum relative distance remained above the specified lower bound ($\alpha = 0.75$; dotted line) for all models, confirming that the van der Waals repulsion restraint worked as intended. Under native guidance, the lower end of the distribution was higher than that with RGI. This is consistent with the higher lower-bound threshold used in native van der Waals guidance (0.775, compared with the RGI default of 0.75).

#### Detailed results of the conformational state-control benchmark

In the QBP benchmark with the interdomain distance restraint, the target distance was increased stepwise from 25.00 Å (closed reference) to 29.10 Å (open reference). In all six models, increasing the target distance progressively decreased the TM-score relative to the closed reference and increased the TM-score relative to the open reference (Fig. S6). The restraint was well satisfied, with residuals close to zero (Fig. S7). In the baseline predictions, the distribution on the two-dimensional TM-score plane depended on the model (Fig. S13). MSA subsampling broadened the distribution of the predictions, but the sampled region remained biased (Fig. S14). In contrast, under the distance-restraint condition, varying the target interdomain distance progressively shifted the predictions from the closed state toward the open state. MolProbity clashscores under the distance-restraint condition were comparable to those under the baseline condition (Fig. S15), although they tended to be slightly higher for Protenix-v2. Thus, in this benchmark, the distance restraint changed the interdomain arrangement while generally maintaining steric clash levels comparable to those in the unrestrained baseline predictions.

With the interdomain angle restraint on ADK, varying the target value of the angle defined by the geometric centroids of the NMP, CORE, and LID domains from 60.9° to 84.8° yielded a stepwise structural change from closed-like to open-like conformations for all six models (Fig. S8). To assess how accurately the restraint was satisfied, the angle was recalculated from the output mmCIF structures using the same atom selections; the maximum absolute deviation from the target value over all 270 structures was 1.33° (Fig. S9). This result indicates that the target angle was achieved with high accuracy not only within the minimization loop but also in the final output structures. For ADK, some models generated structures in the region of TM-score space intermediate between the open and closed references even under the baseline condition (Fig. S16). MSA subsampling also broadened the distribution of the predictions (Fig. S17). However, these control conditions provide no means of specifying the generated state in terms of a target angle. In the MolProbity evaluation, clashscores under the angle-restraint condition were comparable to those under the control conditions for most models, including AlphaFold3 and Boltz-2, whereas a slight increase was observed for ESMFold2 (Fig. S15).

With the RMSD restraints to the two QBP reference structures, the proximity to the two references was controlled in a complementary manner by setting the target RMSD to the open reference to $t$ and that to the closed reference to $T - t$ ($T = 5.3$ Å). The TM-scores averaged over the nine seeds at each target value ranged across models from 0.957 to 0.972 relative to the open reference at $t = 0$ and from 0.982 to 0.985 relative to the closed reference at $t = T$, indicating that the structures reached at the two endpoints agreed well among models (Fig. S10). As with the QBP distance restraint, the conformational change was sampled continuously. The unrestrained baseline and MSA-subsampling controls for this benchmark are the same as those for the QBP distance-restraint benchmark (Figs. S13 and S14), and the RMSD restraints likewise improved on these control conditions. When the target RMSDs to both references were set to 2.65 Å, the mean TM-scores relative to the two references ranged from 0.761 to 0.903, indicating that intermediate structures were obtained (Fig. S10). As expected, however, the restraint residuals did not reach zero (Fig. S11). The distributions of the MolProbity metrics pooled over all target values and seeds showed trends similar to those for the QBP distance restraint and the other benchmarks, with no substantial deterioration. Thus, although the residuals were nonzero, the quality of the sampled structures was not compromised. For Protenix-v2, the clashscores were lower with the RMSD restraints than with the distance restraint, whereas the other models showed nearly the same trends in the two benchmarks (Fig. S15A, C). AlphaFold3 and Boltz-2 in particular tended to give favorable clashscores with little variation (Fig. S15C).

With the custom restraint on the gate-distance difference in DgoT, varying $\Delta D = D_{in} - D_{out}$, the difference between the helix-group distances on the cytoplasmic side ($D_{in}$) and the periplasmic side ($D_{out}$), from −4.3 Å to 5.9 Å steered the structures from outward-facing-like to inward-facing-like conformations for all six models (Fig. S12). When *ΔD* was recalculated from the output mmCIF structures, the maximum absolute deviation from the target value over all 270 structures was approximately 0.01 Å, showing that the custom control variable was also achieved with high accuracy (Fig. S18). On the two-dimensional TM-score plane, the structures generated with RGI changed continuously from outward-facing to inward-facing as *ΔD* was varied, showing that the transition between the two states was sampled (Fig. S12). The two gate distances changed reciprocally (Fig. S19). This behavior is consistent with the rocker-switch model of alternating access, in which the opening of the gate on one side is accompanied by the closing of the gate on the opposite side. The structures at the two endpoints agreed well with the inward- and outward-facing reference structures (Fig. S19). In contrast, with the baseline predictions and MSA subsampling, the two-dimensional TM-score distributions remained biased (Figs. S20 and S21). In the MolProbity evaluation, Protenix-v2 showed a broad distribution of bond-length RMSDs, and the clashscore increased under the custom-restraint condition for some models (Fig. S15); otherwise, no problems were observed. Given cases such as that of Protenix-v2, the results obtained with custom restraints should be evaluated by jointly considering the accuracy with which the control variable is achieved, the similarity to the reference structures, the individual gate conformations, and the quality of the local geometry.

### Supplementary Figure Legends

#### Figure S1

Results for the 209-entry ligand conformer benchmark set.

For each model, baseline (unrestrained) predictions are compared with predictions generated using RGI-Toolkit conformer restraints (RGI). For Boltz-2 and Protenix-v2, predictions obtained with the ligand geometry guidance native to each implementation (Native) are also shown. (A) Fraction of entries in which the chirality of all stereocenters agreed with the input molecule. (B) Fraction of entries in which all distinguishable cis/trans configurations agreed with the input molecule. (C) Bond-length RMSD relative to the ideal conformer. (D) Bond-angle RMSD relative to the ideal conformer. (E) Minimum relative distance between protein and ligand atoms. (F) Ligand pose RMSD. In C–F, box plots show the 10th and 90th percentiles as whiskers, the 25th and 75th percentiles as box boundaries, and the median as a thick black line.

#### Figure S2

Target and achieved ligand bond lengths.

The RGI target bond length (horizontal axis) is plotted against the corresponding bond length in the final predicted structure (vertical axis) for (A) AlphaFold3, (B) Boltz-2, (C) Protenix-v2, (D) Chai-1, (E) OpenFold3, and (F) ESMFold2. Baseline, native guidance (where available), and RGI predictions are shown in gray, orange, and blue, respectively. Each point corresponds to one bond in one predicted ligand. Note that for the baseline, the plotted target values serve only as a reference and were not imposed as restraints. The vertical axis range is fixed, identical to that of the horizontal axis, and shared across all panels; points outside this range are clipped and not shown.

#### Figure S3

Target and achieved ligand bond angles.

The RGI target bond angle is plotted against the corresponding bond angle in the final predicted structure. Each point corresponds to one bond angle, defined by three consecutively bonded atoms, in one predicted ligand. Other details are as in Fig. S2.

#### Figure S4

Target and achieved ligand chiral volumes.

The RGI target signed chiral volume is plotted against the corresponding signed chiral volume in the final predicted structure. Each point corresponds to one tetrahedral stereocenter in one predicted ligand. Because the sign of the chiral volume encodes the handedness of the stereocenter, points in the second and fourth quadrants indicate stereocenters whose chirality is inverted relative to the target. Other details are as in Fig. S2.

#### Figure S5

Target and achieved ligand cis/trans dihedral angles.

The RGI target dihedral angle is plotted against the corresponding dihedral angle in the final predicted structure. Each point corresponds to one double bond with a specified cis/trans configuration in one predicted ligand, with the dihedral angle defined by the four atoms involved in the double bond. Target values near 0° and ±180° correspond to the cis and trans configurations, respectively. Because the dihedral angle is periodic, −180° and 180° are equivalent. Other details are as in Fig. S2.

#### Figure S6

Control of QBP conformational states using an interdomain distance restraint.

(A–C) Representative AlphaFold3 predictions corresponding to the points labeled A–C in panel D, showing closed-like (A), intermediate (B), and open-like (C) conformations. The two domains are shown in dark and light shades. (D–I) Results for AlphaFold3 (D), Boltz-2 (E), Protenix-v2 (F), Chai-1 (G), OpenFold3 (H), and ESMFold2 (I). The target distance between the geometric centroids of the two domains was set to five evenly spaced values from 25.00 to 29.10 Å, and predictions from nine random seeds are shown for each model. The horizontal and vertical axes show the TM-scores against the open (PDB ID: 1GGG) and closed (PDB ID: 1WDN) reference structures, respectively, and point color denotes the target distance. Red and blue stars indicate the open and closed reference structures, respectively.

#### Figure S7

Target and achieved interdomain distances in the QBP distance-restraint benchmark.

The target distance between the geometric centroids of the two domains (horizontal axis) is plotted against the corresponding distance calculated from the final predicted structure (vertical axis) for each model. Each point corresponds to one prediction; 45 predictions are shown per model (five target values with nine random seeds each). The achieved distances closely matched the target values in all models, indicating that the distance restraint was almost fully satisfied.

#### Figure S8

Control of ADK conformational states using a three-group angle restraint.

The target angle defined by the geometric centroids of the NMP, CORE, and LID domains was set to five evenly spaced values from 60.90° to 84.80°, and point color denotes the target angle. The open and closed reference structures are PDB IDs 4AKE and 1AKE, respectively. Even at the target angle of 84.80°, the predictions did not coincide with the open crystal structure, because the angle alone does not uniquely determine the domain arrangement: when the domains twist relative to one another, the angle can reach 84.80° without reproducing the open structure. Panel layout, axes, and symbols are as in Fig. S6.

#### Figure S9

Target and achieved interdomain angles in the ADK angle-restraint benchmark.

The target angle defined by the geometric centroids of the NMP, CORE, and LID domains (horizontal axis) is plotted against the corresponding angle calculated from the final predicted structure (vertical axis) for each model. Each point corresponds to one prediction; 45 predictions are shown per model (five target values with nine random seeds each). The achieved angles closely matched the target values in all models, indicating that the angle restraint was well satisfied

**Figure S10**

Control of QBP conformational states using dual RMSD restraints against the open and closed references.

The target RMSDs against the open (PDB ID: 1GGG) and closed (PDB ID: 1WDN) references were set to $t$ and $T - t$, respectively, where $T = 5.3$ Å is the RMSD between the two references, and $t$ was set to five evenly spaced values from 0 to 5.3 Å. Point color denotes $t$. Panel layout, axes, and symbols are as in Fig. S6.

**Figure S11**

Target and achieved Cα RMSDs in the QBP RMSD-restraint benchmark.

The target Cα RMSD (horizontal axis) is plotted against the corresponding Cα RMSD calculated from the final predicted structure (vertical axis) for each model. Two points are shown for each prediction, one for the RMSD to the open reference (1GGG; target $t$) and the other for the RMSD to the closed reference (1WDN; target $T - t$), shown in blue circles and orange triangles, respectively. Predictions from five values of $t$ with nine random seeds each are shown per model. The achieved RMSDs did not exactly match the target values. These residual deviations are expected from the restraint design rather than a failure of the restraints (see main text).

#### Figure S12

Control of DgoT conformational states using a custom restraint defined by the difference between the cytoplasmic and periplasmic gate distances.

The control variable was defined as $\Delta D = D_{in} - D_{out}$, where $D_{in}$ and $D_{out}$ are the distances between the geometric centroids of the transmembrane helix groups on the cytoplasmic and periplasmic sides, respectively, and its target value was set to five evenly spaced values from −4.30 to 5.90 Å. The horizontal and vertical axes show the TM-scores against the outward-facing (PDB ID: 6E9O; red star) and inward-facing (PDB ID: 6E9N; blue star) reference structures, respectively, and point color denotes the target $\Delta D$. Panel layout and other details are as in Fig. S6.

#### Figure S13

Two-dimensional TM-score distributions of QBP predictions under the baseline condition.

Predictions generated with the full MSA, without restraints or MSA subsampling, are shown on the same two-dimensional TM-score plane as in Fig. S6. Some models yielded both open-like and closed-like structures, but no intermediate structures were obtained. Other details are as in Fig. S6.

#### Figure S14

Two-dimensional TM-score distributions of QBP predictions obtained with MSA subsampling.

Predictions generated with subsampled MSAs are shown on the same two-dimensional TM-score plane as in Fig. S6, with point color denoting the MSA depth (16, 32, 64, 128, or 256). MSA subsampling sampled a broader range of conformations than the baseline, but the sampled region remained biased. Other details are as in Fig. S6.

#### Figure S15

MolProbity stereochemical quality in the baseline, MSA-subsampling, and RGI conditions across six models.

Predicted structures from the state-control benchmarks were evaluated with MolProbity: (A) QBP with the interdomain distance restraint, (B) ADK with the interdomain angle restraint, (C) QBP with the dual-reference RMSD restraints, and (D) DgoT with the custom restraint. In each row, the panels show, from left to right, bond-angle RMSD, bond-length RMSD, and MolProbity clashscore; lower values are better for all three metrics. Within each model, gray, orange, and blue boxes denote the baseline, MSA-subsampling, and RGI conditions, respectively. The baseline and MSA-subsampling distributions in C are the same QBP control predictions as those shown in A. Box plots show the 10th and 90th percentiles as whiskers, the 25th and 75th percentiles as box boundaries, and the median as a thick black line. For each metric, the same y-axis range is used in A–D.

#### Figure S16

Two-dimensional TM-score distributions of ADK predictions under the baseline condition.

Predictions generated with the full MSA, without restraints or MSA subsampling, are shown on the same two-dimensional TM-score plane as in Fig. S8. Intermediate structures tended to be sampled even under this condition, but the sampled region remained biased. Unlike the other models, Protenix-v2 sampled the open state. Other details are as in Fig. S8.

#### Figure S17

Two-dimensional TM-score distributions of ADK predictions obtained with MSA subsampling.

Predictions generated with subsampled MSAs are shown on the same two-dimensional TM-score plane as in Fig. S8, with point color denoting the MSA depth (16, 32, 64, 128, or 256). MSA subsampling enhanced sampling in some models, such as OpenFold3; however, the overall distributions differed little from those obtained under the baseline condition. Other details are as in Fig. S8.

#### Figure S18

Target and achieved $\Delta D$ in the DgoT custom-restraint benchmark.

The target value of $\Delta D = D_{\text{in}} - D_{\text{out}}$ (horizontal axis) is plotted against the corresponding value calculated from the final predicted structure (vertical axis) for each model. Each point corresponds to one prediction; 45 predictions are shown per model (five target values with nine random seeds each). The achieved values closely matched the target values in all models, indicating that the custom restraint was well satisfied.

#### Figure S19

Periplasmic and cytoplasmic gate distances in DgoT predictions generated with the custom restraint.

For each model, the periplasmic gate distance Dout (horizontal axis) and the cytoplasmic gate distance Din (vertical axis) are plotted, where Dout and Din are the distances between the geometric centroids of the periplasmic helix groups (C and D) and the cytoplasmic helix groups (A and B), respectively. Both axes span 15–30 Å. The reference structures are shown as a red star for the outward-facing state (6E9O) and a blue star for the inward-facing state (6E9N). Point color denotes the target value of $\Delta D$. As $\Delta D$ is swept from negative to positive values, the two gate distances change reciprocally in all six models: the periplasmic gate opens as the cytoplasmic gate closes at $\Delta D < 0$ (outward-facing), and the reverse occurs at $\Delta D > 0$ (inward-facing). This behavior is consistent with the rocker-switch model of alternating access. The predictions at the two ends of the sweep agreed well with the outward-facing and inward-facing reference structures.

#### Figure S20

Two-dimensional TM-score distributions of DgoT predictions under the baseline condition.

Predictions generated with the full MSA, without restraints or MSA subsampling, are shown on the same two-dimensional TM-score plane as in Fig. S12. The distribution of the predictions was biased, and the transition between the outward-facing and inward-facing states was not sampled. Other details are as in Fig. S12.

#### Figure S21

Two-dimensional TM-score distributions of DgoT predictions obtained with MSA subsampling.

Predictions generated with subsampled MSAs are shown on the same two-dimensional TM-score plane as in Fig. S12, with point color denoting the MSA depth (16, 32, 64, 128, or 256). With MSA subsampling, only Protenix-v2 tended to sample both the outward-facing and inward-facing states, including intermediate structures between them. For the other models, the distribution of the predictions remained biased, in contrast to the continuous transition between the two states obtained with the custom restraint (Fig. S12). Other details are as in Fig. S12.

(11) Leano, J. B.; Batarni, S.; Eriksen, J.; Juge, N.; Pak, J. E.; Kimura-Someya, T.; Robles-Colmenares, Y.; Moriyama, Y.; Stroud, R. M.; Edwards, R. H. Structures Suggest a Mechanism for Energy Coupling by a Family of Organic Anion

Transporters. *PLoS Biol.* **2019**, *17* (5), e3000260.

(12) Zhang, C.; Shine, M.; Pyle, A. M.; Zhang, Y. US-Align: Universal Structure Alignments of Proteins, Nucleic Acids, and Macromolecular Complexes. *Nat. Methods* **2022**, *19* (9), 1109–1115.

(13) Williams, C. J.; Headd, J. J.; Moriarty, N. W.; Prisant, M. G.; Videau, L. L.; Deis, L. N.; Verma, V.; Keedy, D. A.; Hintze, B. J.; Chen, V. B.; Jain, S.; Lewis, S. M.; Arendall, W. B., 3rd; Snoeyink, J.; Adams, P. D.; Lovell, S. C.; Richardson, J. S.; Richardson, D. C. MolProbity: More and Better Reference Data for Improved All-Atom Structure Validation. *Protein Sci.* **2018**, *27* (1), 293–315.
