## Supplementary material for "RGI-Toolkit: Differentiable Restraints for Controllable Biomolecular Structure Prediction": suppl_fig

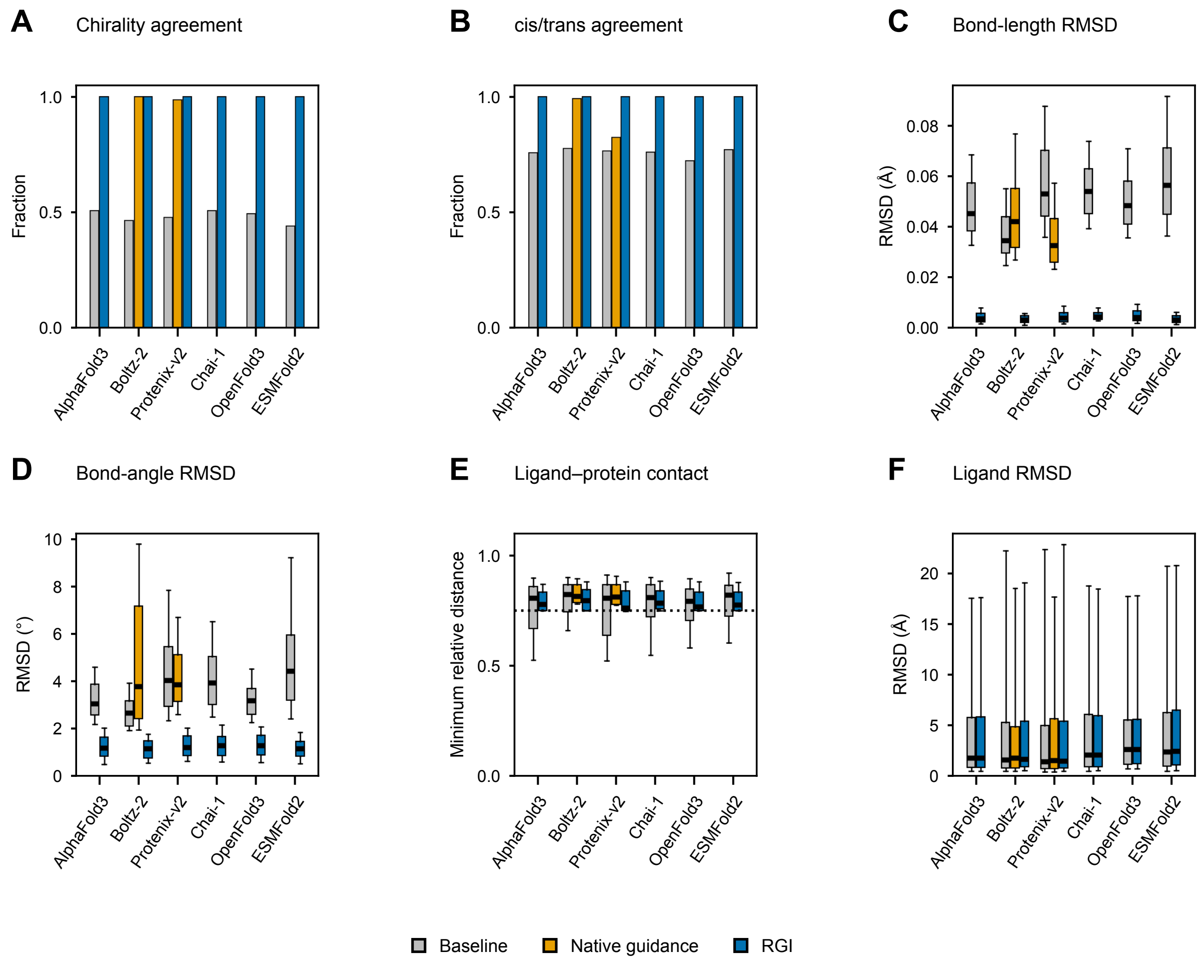
A
Chirality agreement
Fraction
B
cis/trans agreement
Fraction
C
Bond-length RMSD
RMSD (Å)
D
Bond-angle RMSD
RMSD (°)
E
Ligand–protein contact
Minimum relative distance
F
Ligand RMSD
RMSD (Å)
AlphaFold3
Boltz-2
Protenix-v2
Chai-1
OpenFold3
ESMFold2
Baseline
Native guidance
RGI

### Figure S1

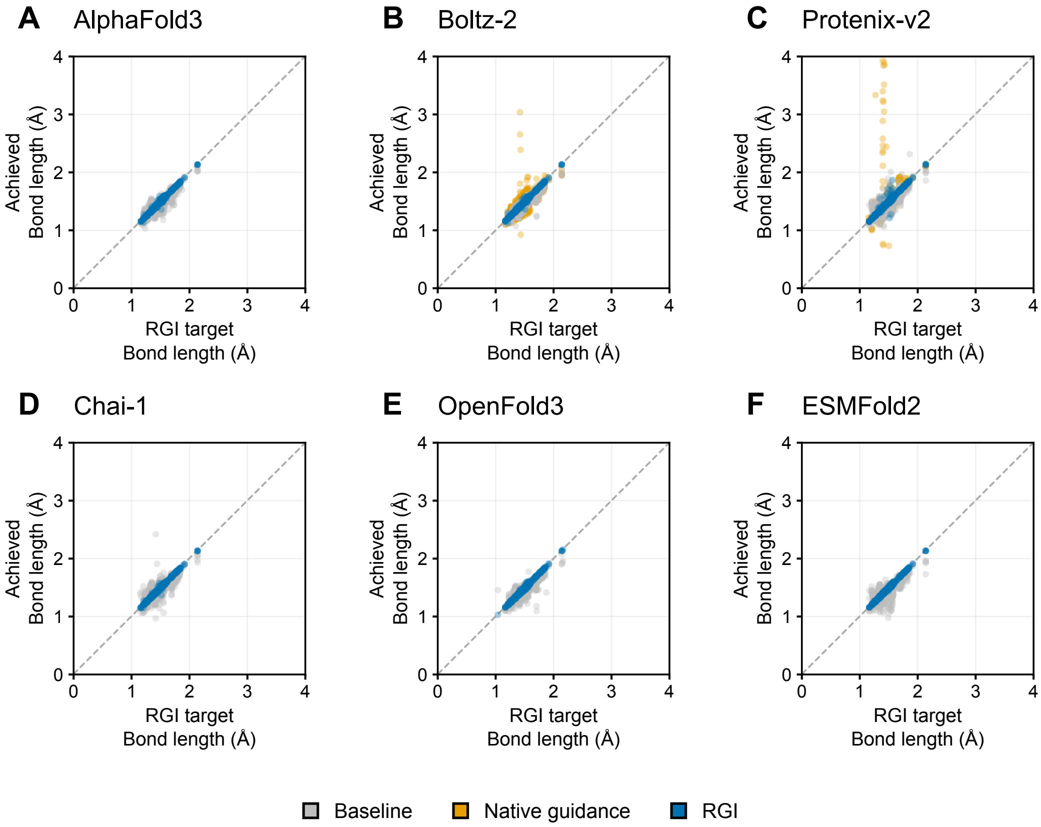
A
AlphaFold3
B
Boltz-2
C
Protenix-v2
D
Chai-1
E
OpenFold3
F
ESMFold2
Achieved
Bond length (Å)
RGI target
Bond length (Å)
0
1
2
3
4
Baseline
Native guidance
RGI

### Figure S2

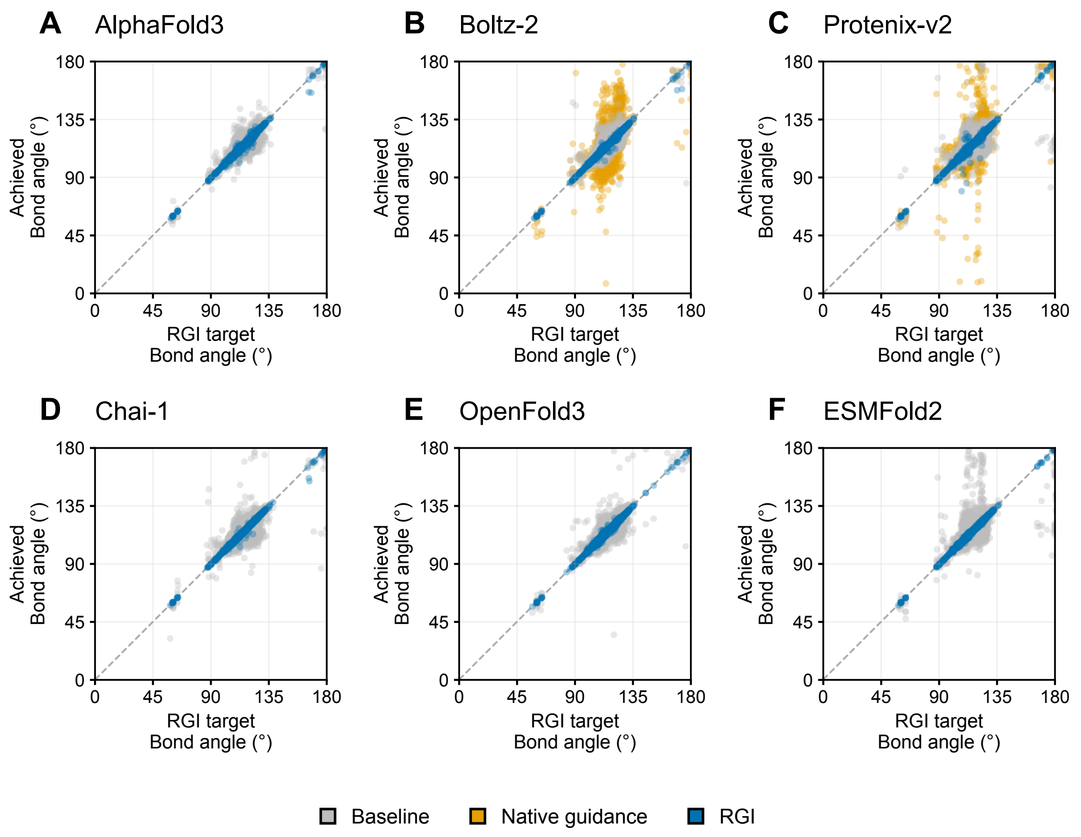
A AlphaFold3
B Boltz-2
C Protenix-v2
D Chai-1
E OpenFold3
F ESMFold2
Achieved Bond angle (°)
RGI target Bond angle (°)
0
45
90
135
180
Baseline
Native guidance
RGI

### Figure S3

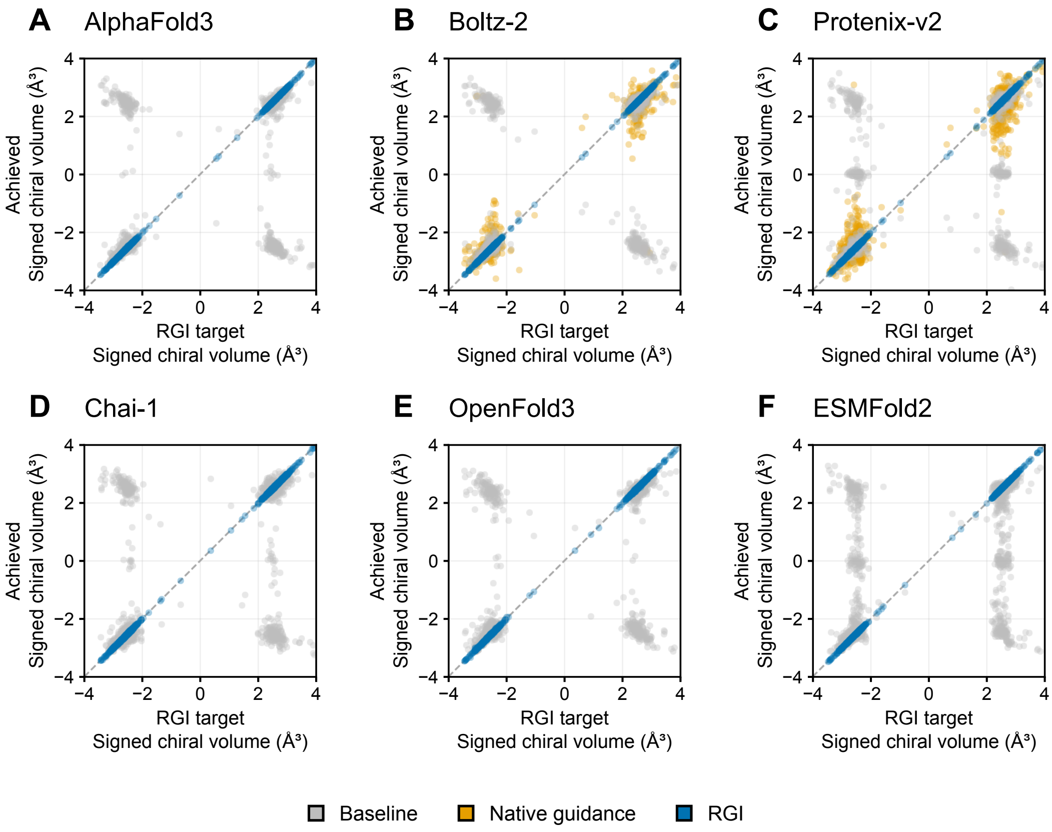

A
AlphaFold3
B
Boltz-2
C
Protenix-v2
D
Chai-1
E
OpenFold3
F
ESMFold2
Achieved
Signed chiral volume (Å³)
RGI target
Signed chiral volume (Å³)
Baseline
Native guidance
RGI


### Figure S4

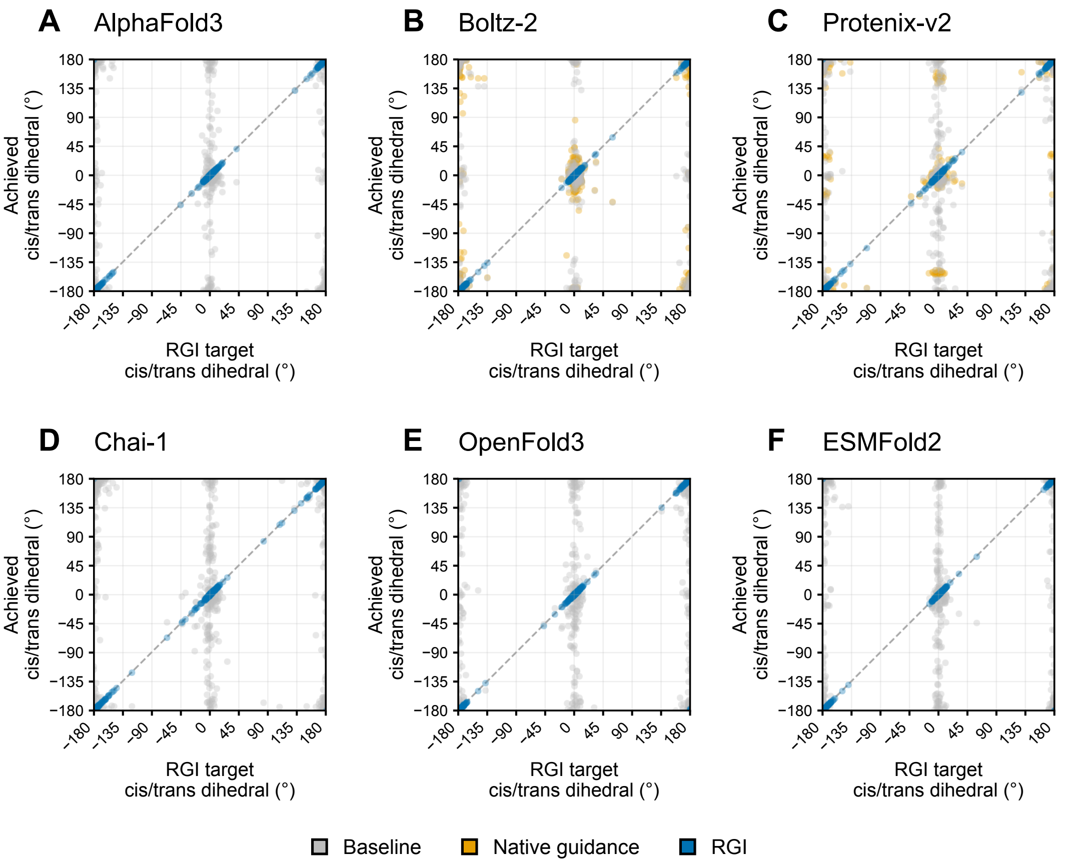

A
AlphaFold3
B
Boltz-2
C
Protenix-v2
D
Chai-1
E
OpenFold3
F
ESMFold2
Achieved
cis/trans dihedral (°)
RGI target
cis/trans dihedral (°)
180
135
90
45
0
−45
−90
−135
−180
Baseline
Native guidance
RGI


### Figure S5

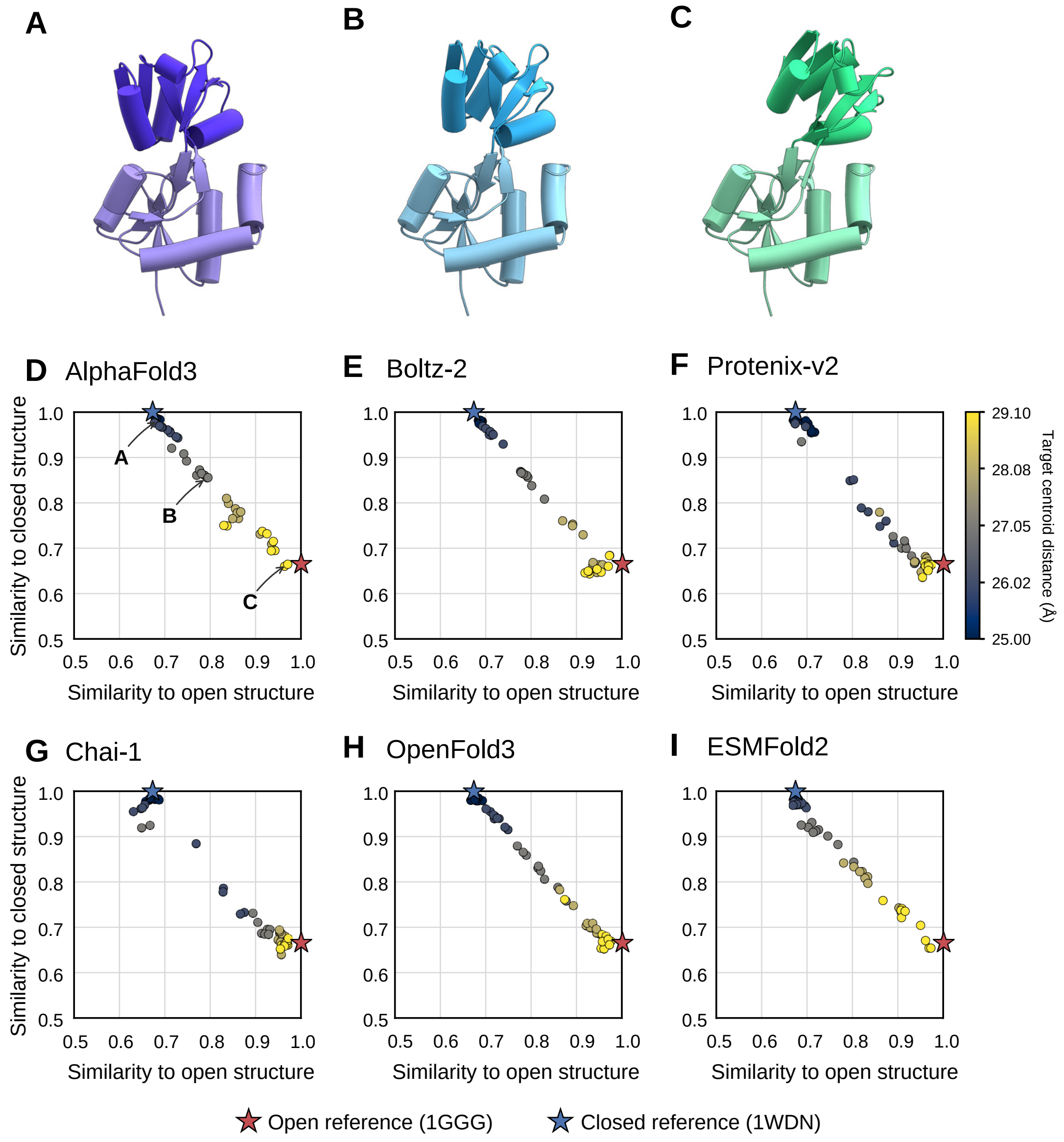
A
B
C
D AlphaFold3
E Boltz-2
F Protenix-v2
G Chai-1
H OpenFold3
I ESMFold2
Similarity to closed structure
Similarity to open structure
1.0
0.9
0.8
0.7
0.6
0.5
29.10
28.08
27.05
26.02
25.00
Target centroid distance (Å)
Open reference (1GGG)
Closed reference (1WDN)

### Figure S6

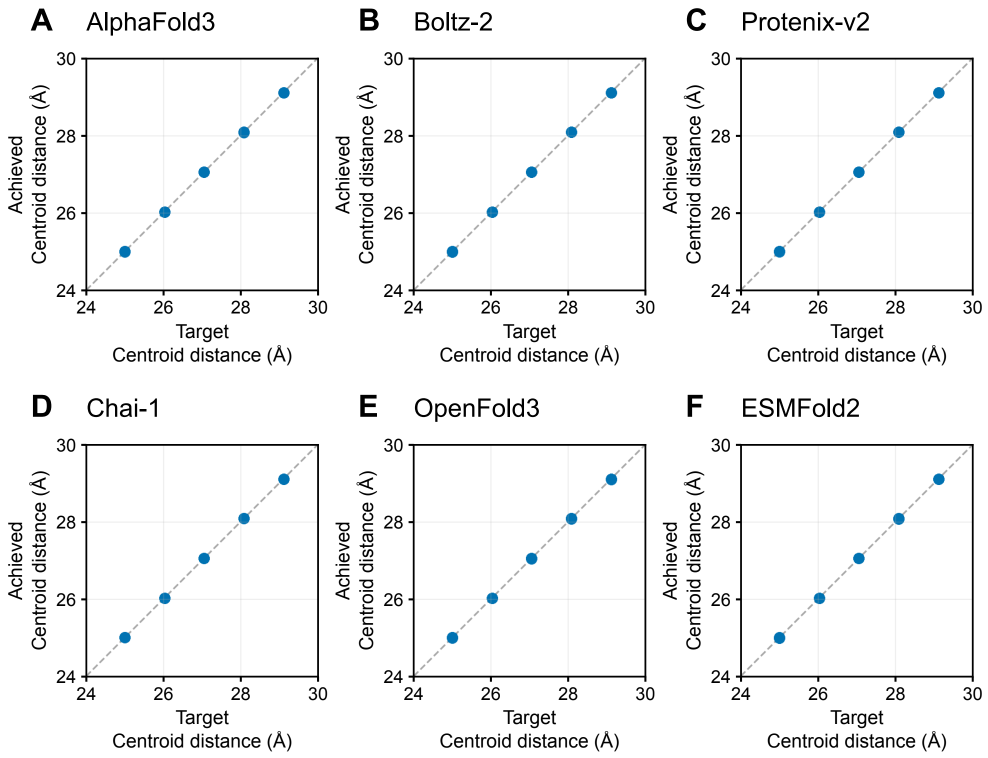
A
AlphaFold3
B
Boltz-2
C
Protenix-v2
D
Chai-1
E
OpenFold3
F
ESMFold2
Achieved
Centroid distance (Å)
Target
Centroid distance (Å)
24
26
28
30

### Figure S7

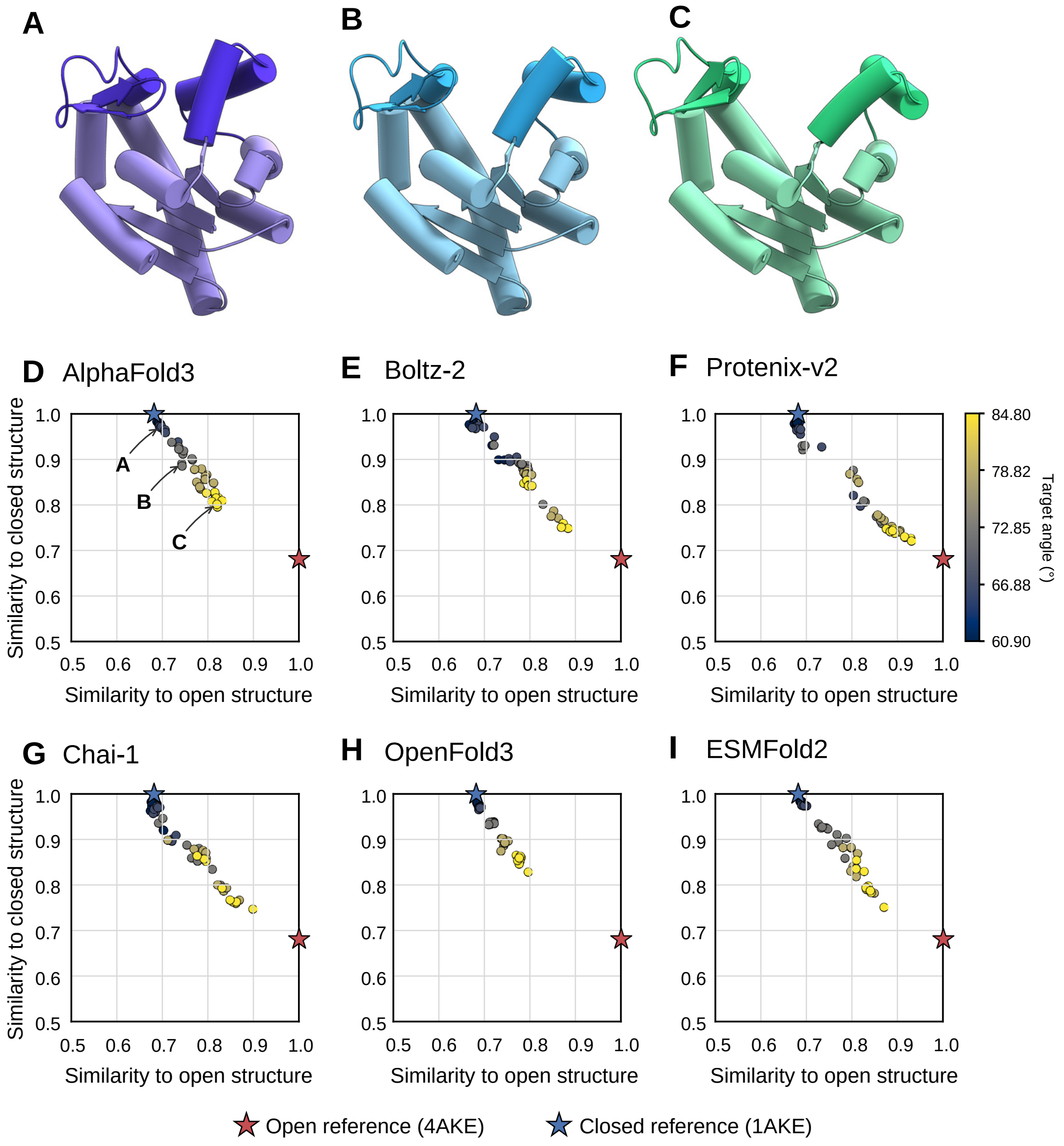
A
B
C
D AlphaFold3
E Boltz-2
F Protenix-v2
G Chai-1
H OpenFold3
I ESMFold2
Similarity to closed structure
Similarity to open structure
Target angle (°)
84.80
78.82
72.85
66.88
60.90
Open reference (4AKE)
Closed reference (1AKE)

### Figure S8

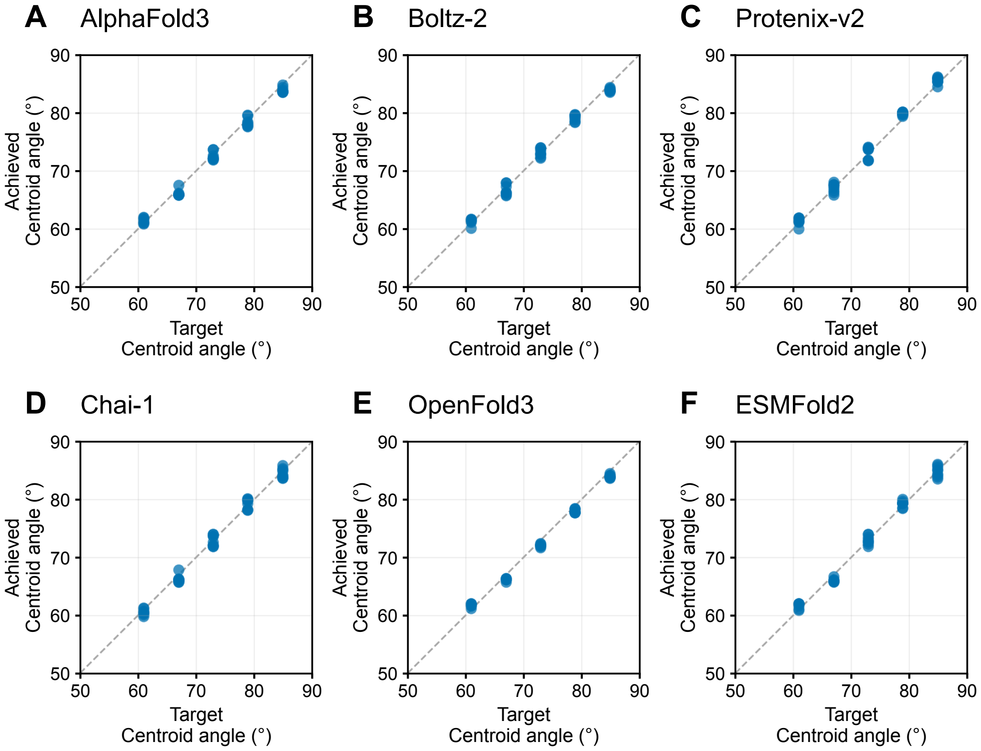
A AlphaFold3
B Boltz-2
C Protenix-v2
D Chai-1
E OpenFold3
F ESMFold2
Achieved Centroid angle (°)
Target Centroid angle (°)
50 60 70 80 90

### Figure S9

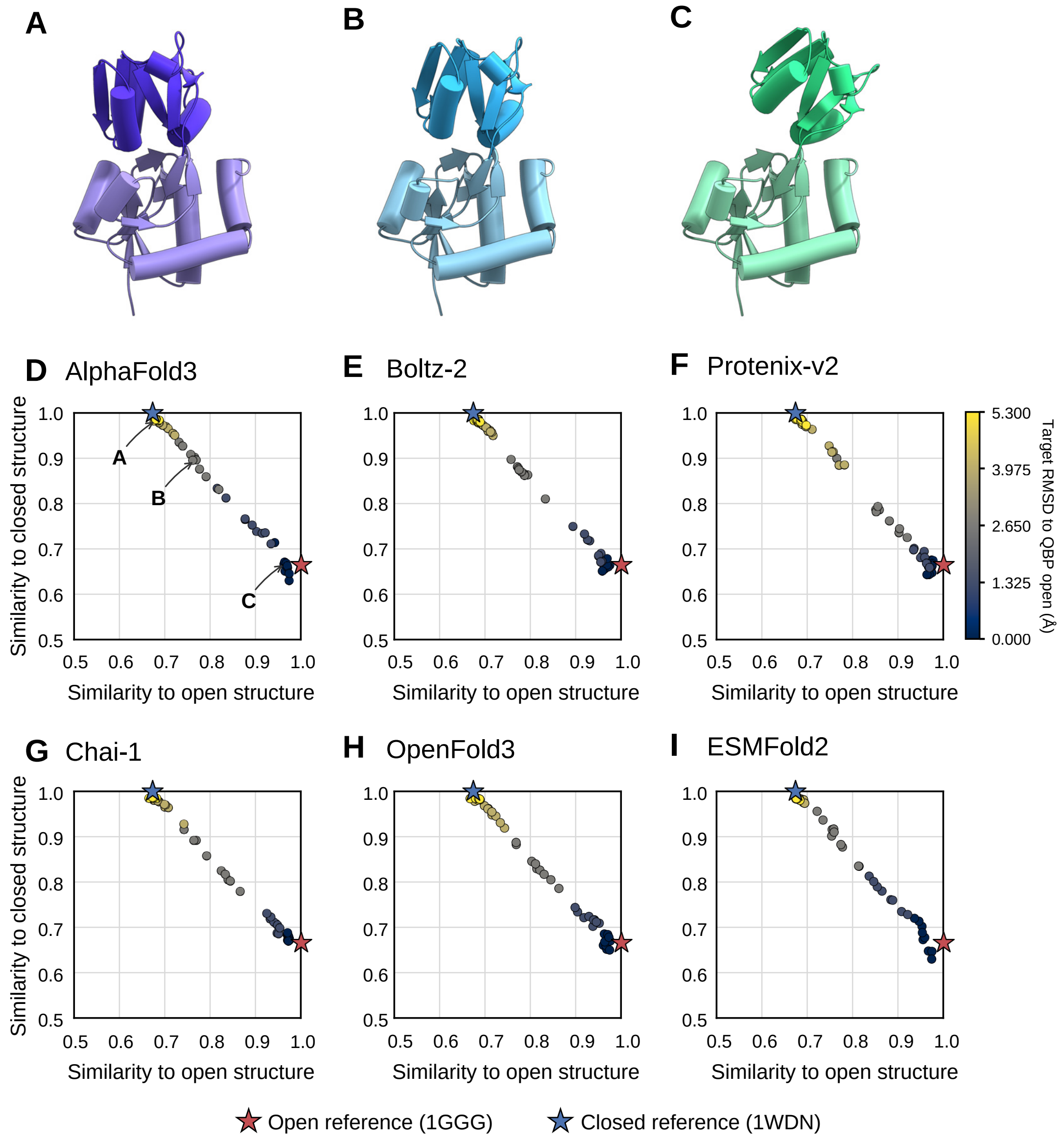
A
B
C
D AlphaFold3
E Boltz-2
F Protenix-v2
G Chai-1
H OpenFold3
I ESMFold2
Similarity to closed structure
Similarity to open structure
Target RMSD to QBP open (Å)
5.300
3.975
2.650
1.325
0.000
Open reference (1GGG)
Closed reference (1WDN)

### Figure S10

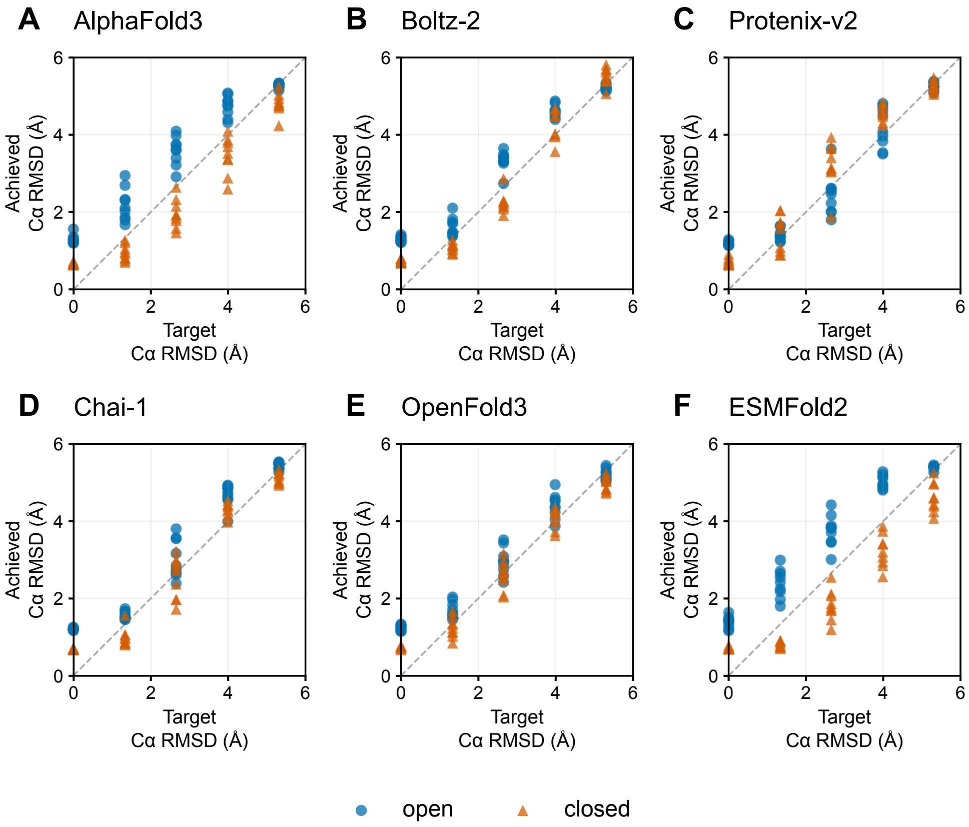

A
AlphaFold3
B
Boltz-2
C
Protenix-v2
D
Chai-1
E
OpenFold3
F
ESMFold2
Achieved
Cα RMSD (Å)
Target
Cα RMSD (Å)
open
closed


### Figure S11

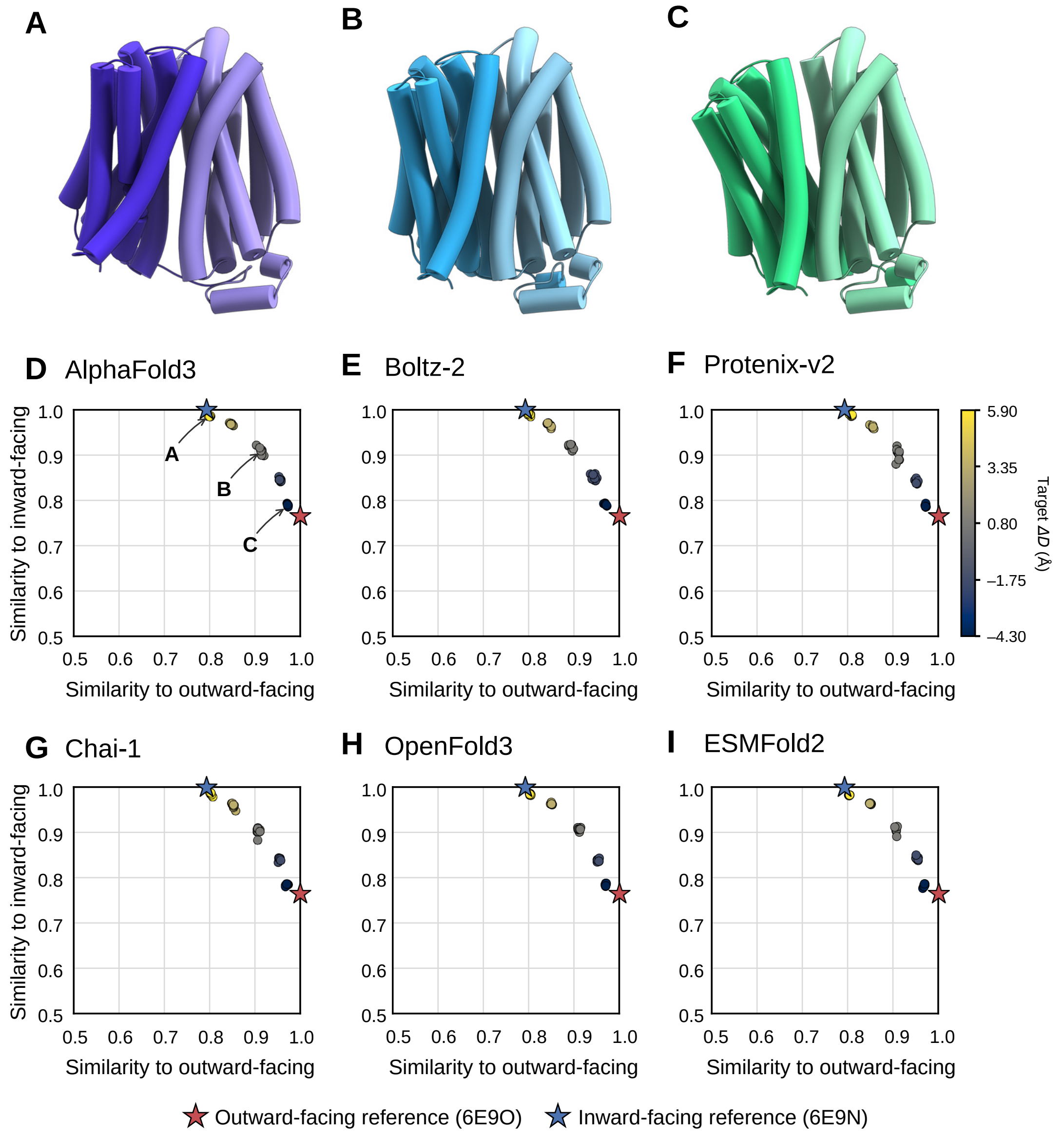

A
B
C
D AlphaFold3
E Boltz-2
F Protenix-v2
G Chai-1
H OpenFold3
I ESMFold2
Similarity to inward-facing
Similarity to outward-facing
Target ΔD (Å)
5.90
3.35
0.80
−1.75
−4.30
Outward-facing reference (6E9O)
Inward-facing reference (6E9N)


### Figure S12

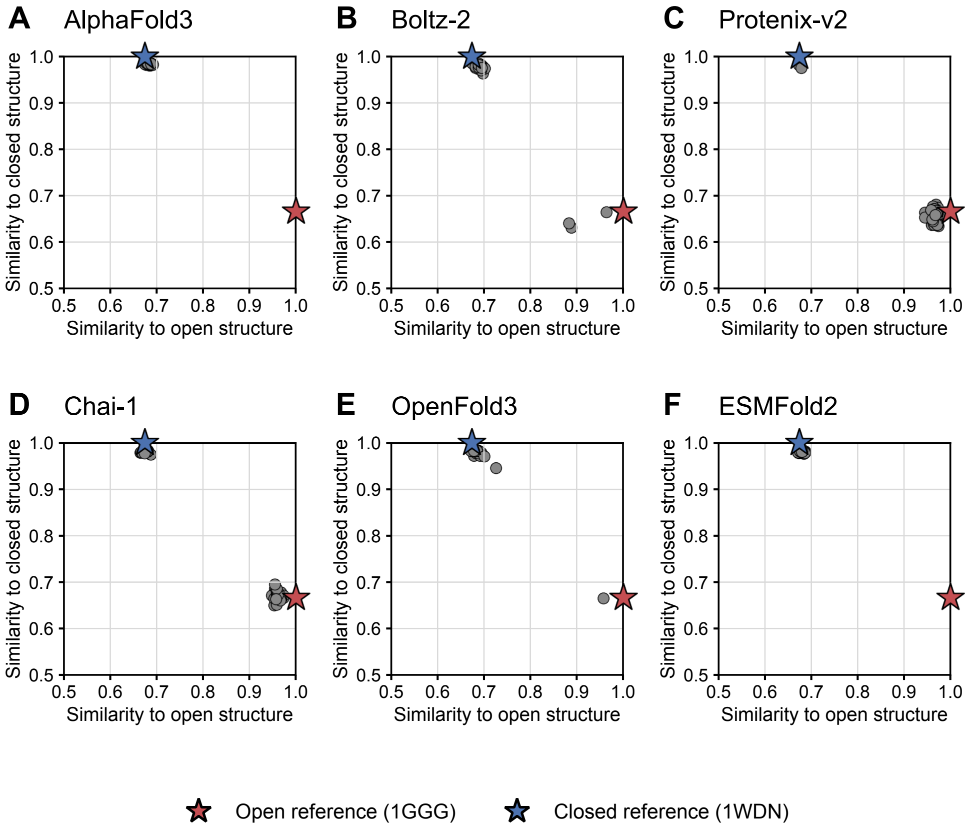

A
AlphaFold3
B
Boltz-2
C
Protenix-v2
D
Chai-1
E
OpenFold3
F
ESMFold2
Similarity to closed structure
Similarity to open structure
0.5
0.6
0.7
0.8
0.9
1.0
Open reference (1GGG)
Closed reference (1WDN)


**Figure S13**

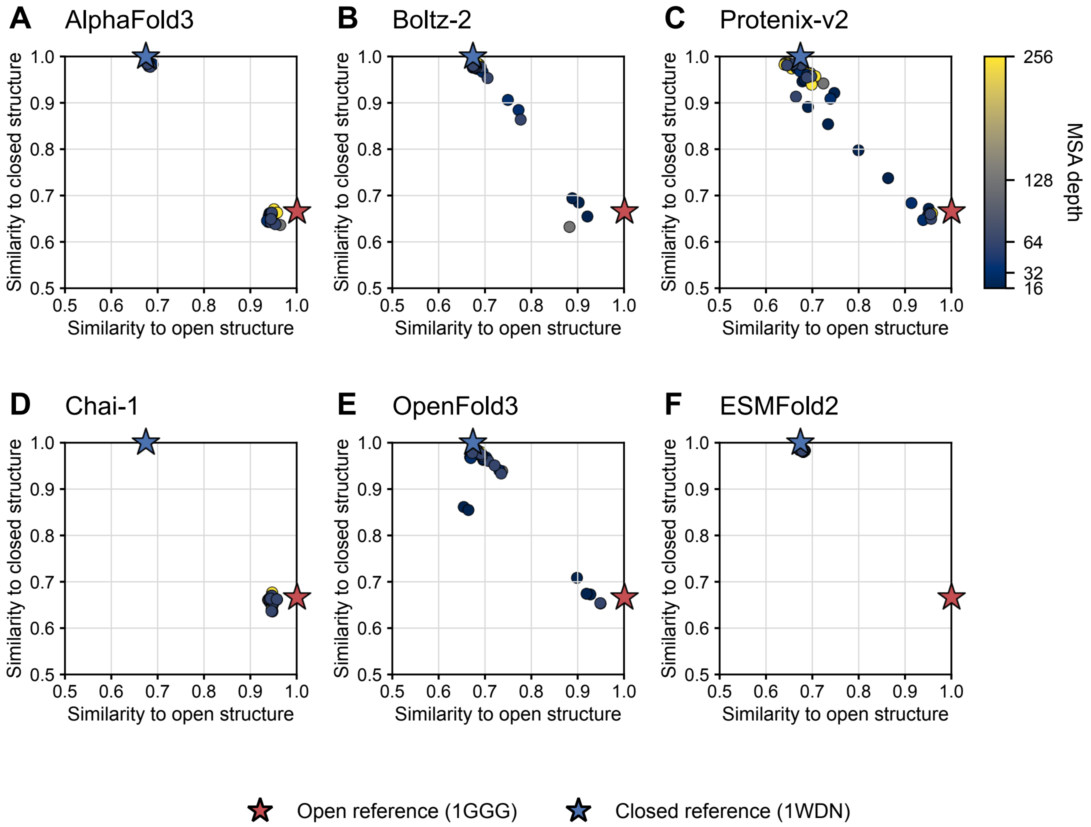
A AlphaFold3
B Boltz-2
C Protenix-v2
D Chai-1
E OpenFold3
F ESMFold2
Similarity to closed structure
Similarity to open structure
0.5
0.6
0.7
0.8
0.9
1.0
MSA depth
256
128
64
32
16
Open reference (1GGG)
Closed reference (1WDN)

**Figure S14**

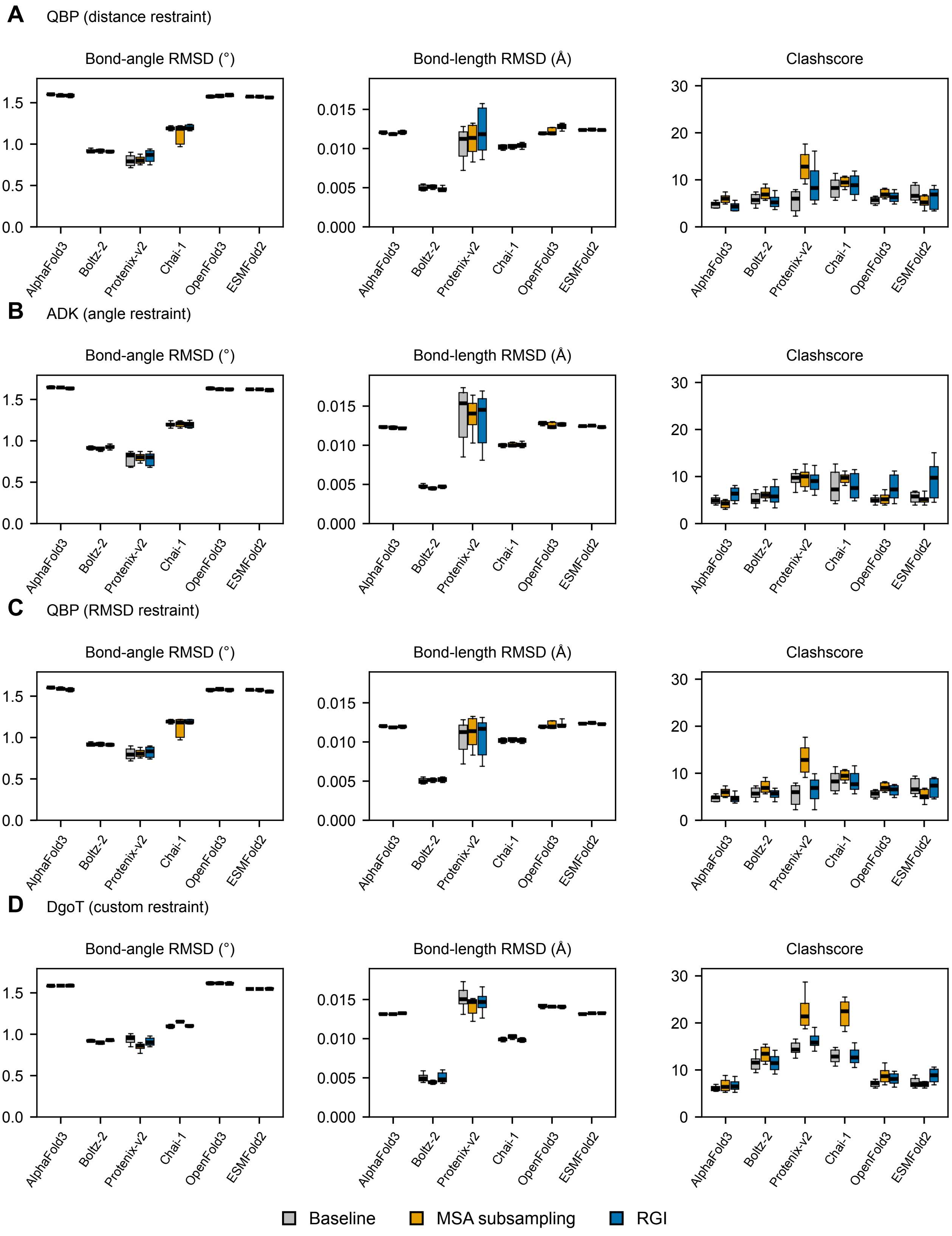
A
QBP (distance restraint)
B
ADK (angle restraint)
C
QBP (RMSD restraint)
D
DgoT (custom restraint)
Bond-angle RMSD (°)
Bond-length RMSD (Å)
Clashscore
AlphaFold3
Boltz-2
Protenix-v2
Chai-1
OpenFold3
ESMFold2
Baseline
MSA subsampling
RGI

### Figure S15

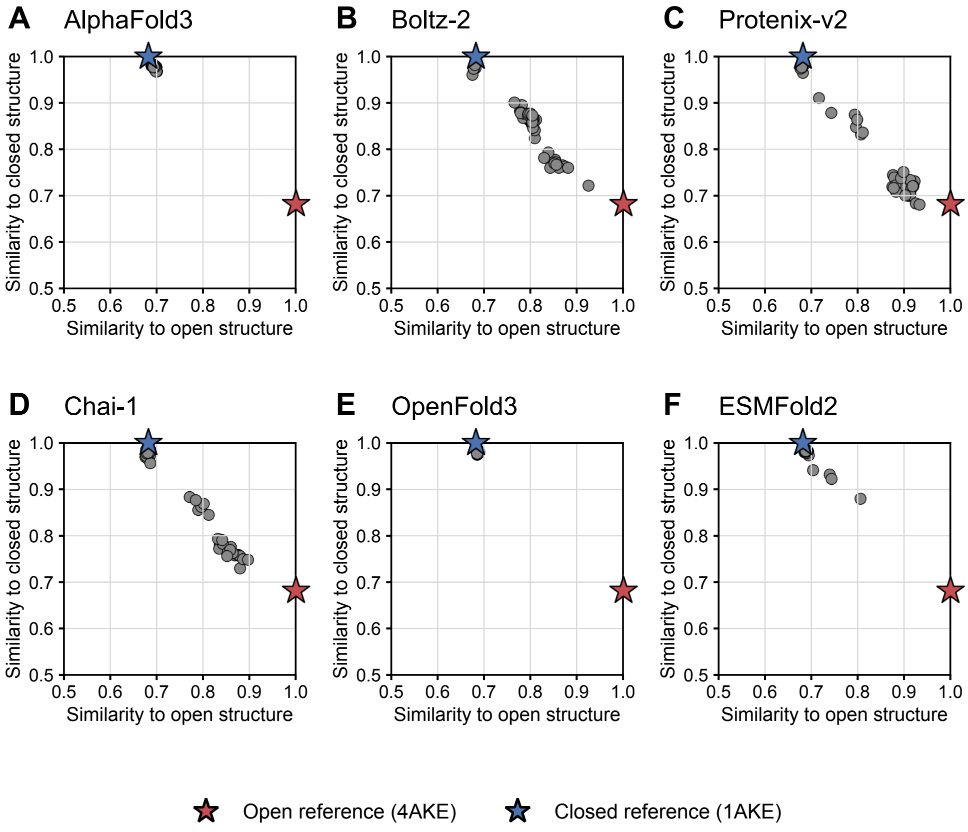
A
AlphaFold3
B
Boltz-2
C
Protenix-v2
D
Chai-1
E
OpenFold3
F
ESMFold2
Similarity to closed structure
Similarity to open structure
0.5
0.6
0.7
0.8
0.9
1.0
Open reference (4AKE)
Closed reference (1AKE)

**Figure S16**

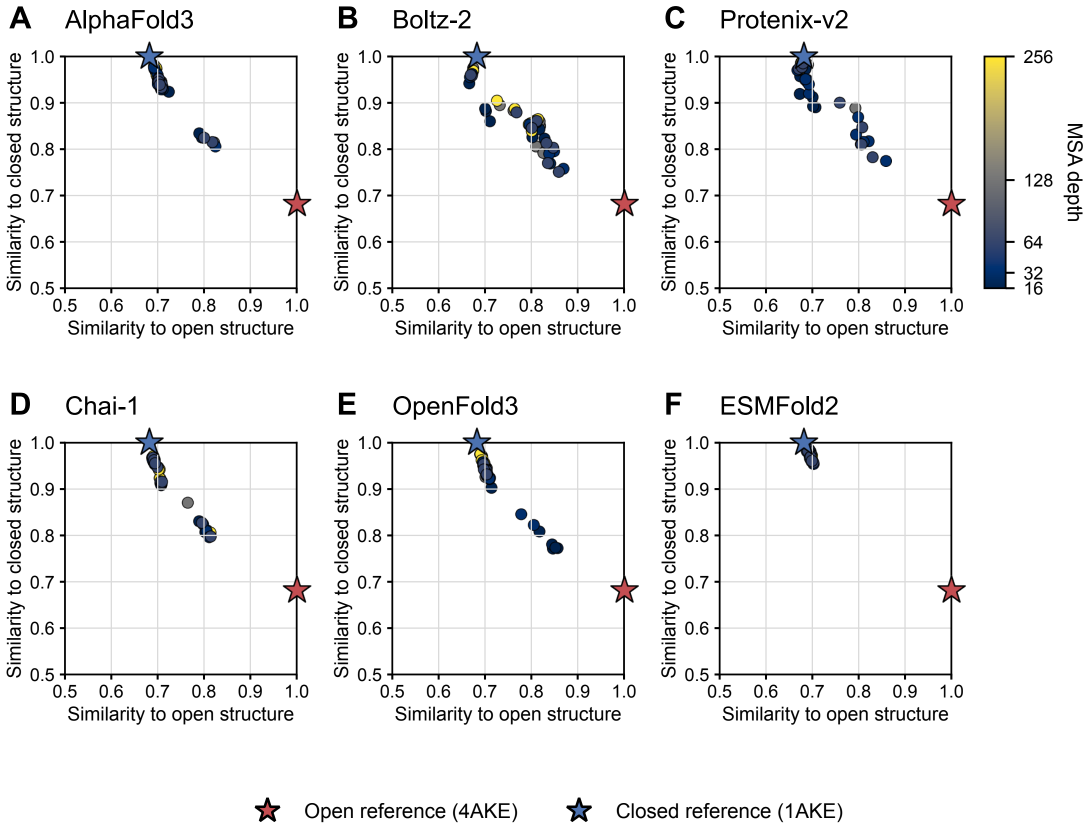
A AlphaFold3
B Boltz-2
C Protenix-v2
D Chai-1
E OpenFold3
F ESMFold2
Similarity to closed structure
Similarity to open structure
MSA depth
256
128
64
32
16
Open reference (4AKE)
Closed reference (1AKE)

**Figure S17**

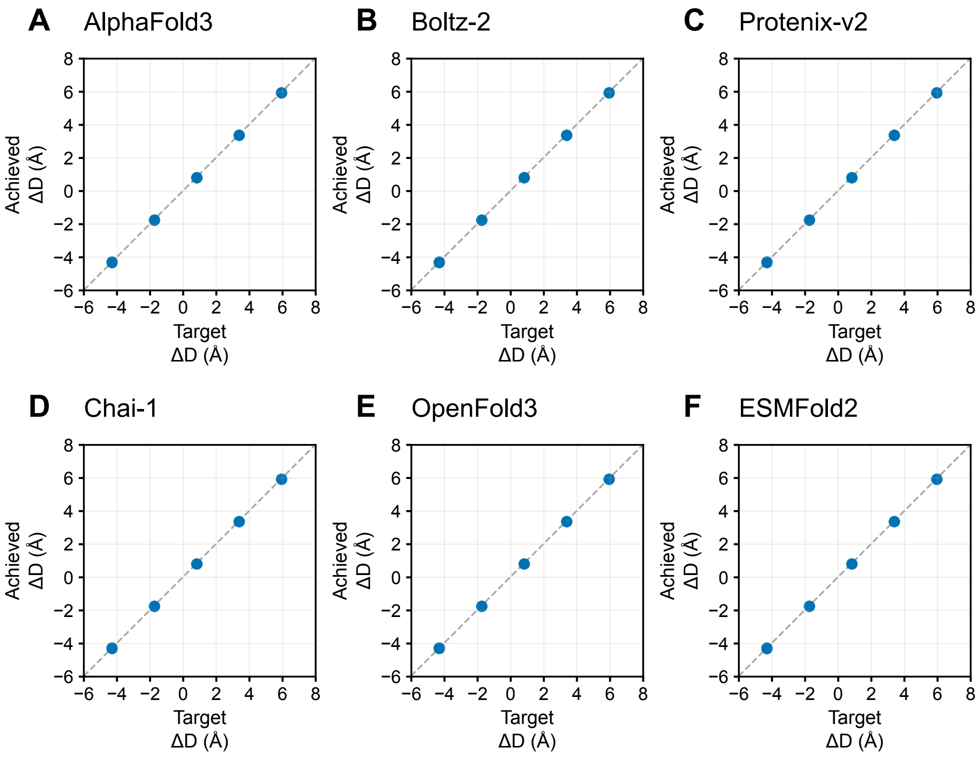

A
AlphaFold3
B
Boltz-2
C
Protenix-v2
D
Chai-1
E
OpenFold3
F
ESMFold2
Achieved
ΔD (Å)
Target
ΔD (Å)


### Figure S18

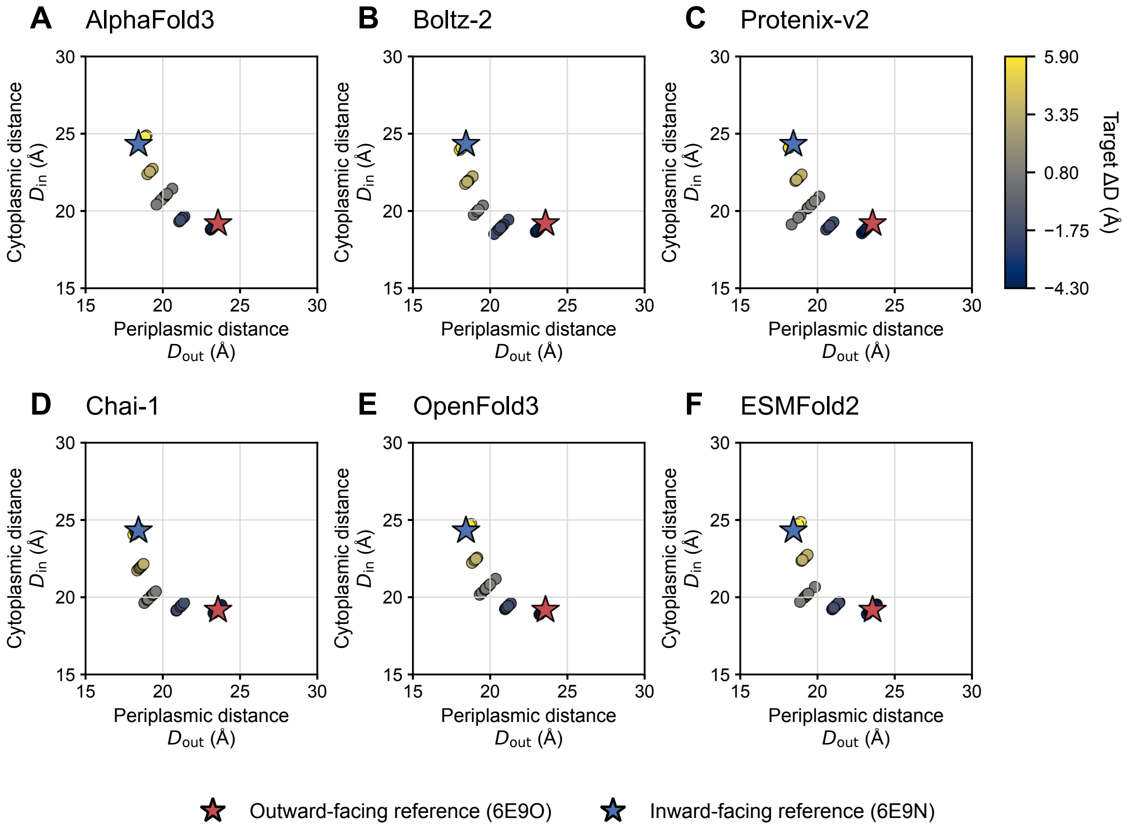

A AlphaFold3
B Boltz-2
C Protenix-v2
D Chai-1
E OpenFold3
F ESMFold2
Cytoplasmic distance $D_{in}$ (Å)
Periplasmic distance $D_{out}$ (Å)
Target ΔD (Å)
5.90
3.35
0.80
−1.75
−4.30
Outward-facing reference (6E9O)
Inward-facing reference (6E9N)


### Figure S19

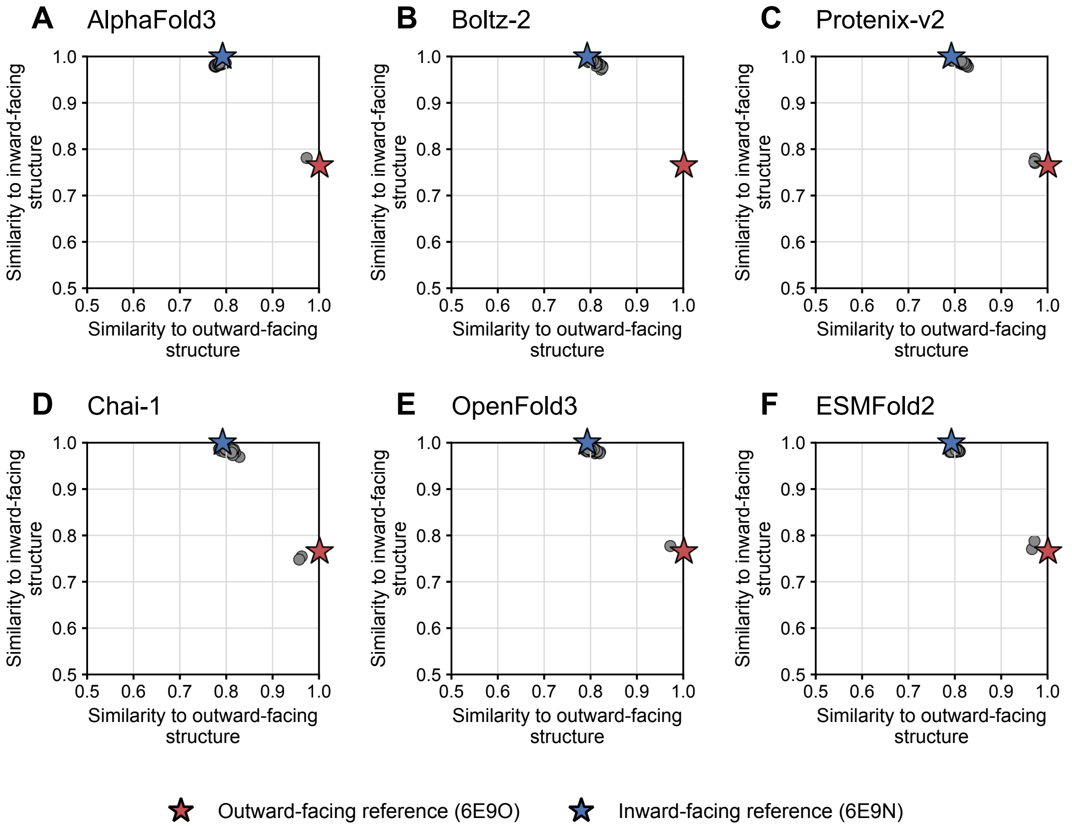
A
AlphaFold3
B
Boltz-2
C
Protenix-v2
D
Chai-1
E
OpenFold3
F
ESMFold2
Similarity to inward-facing structure
Similarity to outward-facing structure
0.5
0.6
0.7
0.8
0.9
1.0
Outward-facing reference (6E9O)
Inward-facing reference (6E9N)

### Figure S20

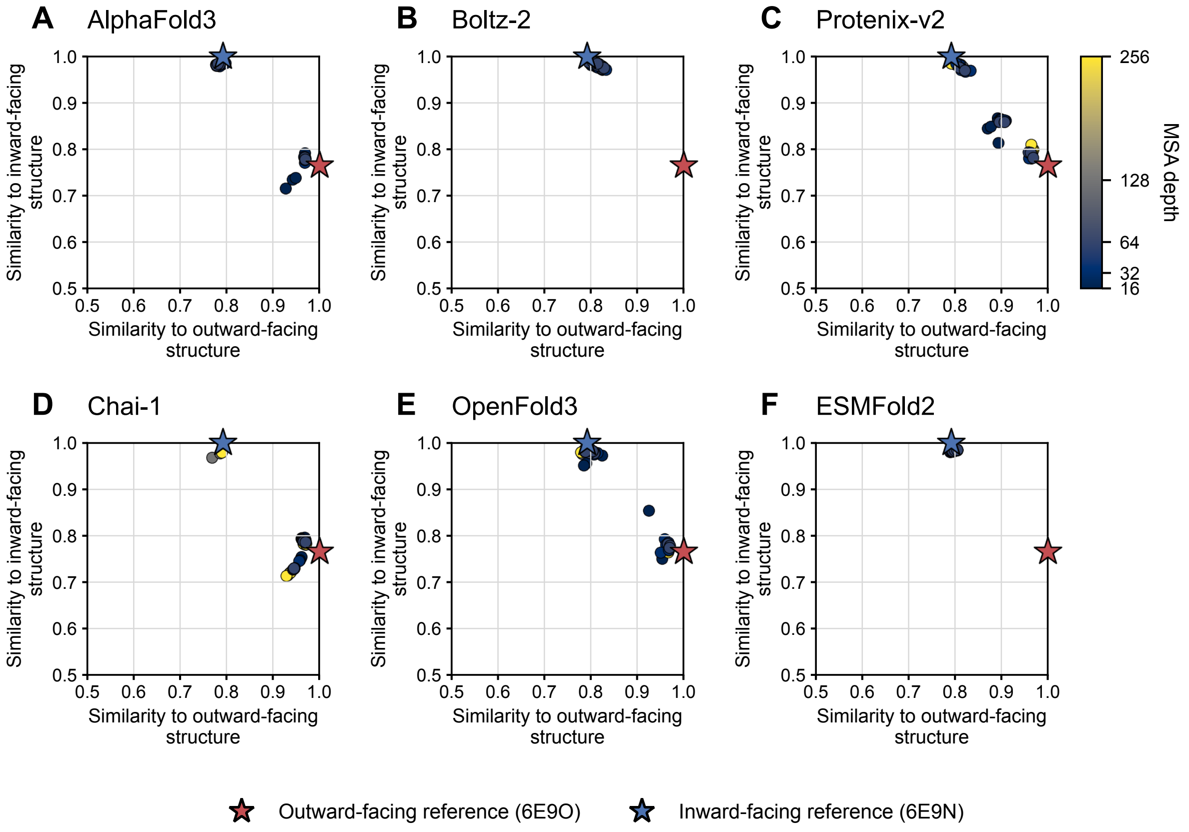
A AlphaFold3
B Boltz-2
C Protenix-v2
D Chai-1
E OpenFold3
F ESMFold2
Similarity to inward-facing structure
Similarity to outward-facing structure
0.5 0.6 0.7 0.8 0.9 1.0
MSA depth
256
128
64
32
16
Outward-facing reference (6E9O)
Inward-facing reference (6E9N)

**Figure S21**
